# Molecular Signatures of Resilience to Alzheimer’s Disease in Neocortical Layer 4 Neurons

**DOI:** 10.1101/2024.11.03.621787

**Authors:** S Akila Parvathy Dharshini, Jorge Sanz-Ros, Jie Pan, Weijing Tang, Kristen Vallejo, Marcos Otero-Garcia, Inma Cobos

## Abstract

Single-cell omics is advancing our understanding of selective neuronal vulnerability in Alzheimer’s disease (AD), revealing specific subtypes that are either susceptible or resilient to neurodegeneration. Using single-nucleus and spatial transcriptomics to compare neocortical regions affected early (prefrontal cortex and precuneus) or late (primary visual cortex) in AD, we identified a resilient excitatory population in layer 4 of the primary visual cortex expressing *RORB*, *CUX2*, and *EYA4*. Layer 4 neurons in association neocortex also remained relatively preserved as AD progressed and shared overlapping molecular signatures of resilience. Early in the disease, resilient neurons upregulated genes associated with synapse maintenance, synaptic plasticity, calcium homeostasis, and neuroprotective factors, including *GRIN2A, RORA, NRXN1, NLGN1, NCAM2, FGF14, NRG3, NEGR1*, and *CSMD1*. We also identified *KCNIP4*, which encodes a voltage-gated potassium (Kv) channel-interacting protein that interacts with Kv4.2 channels and presenilins, as a key factor linked to resilience. *KCNIP4* was consistently upregulated in the early stages of pathology. Furthermore, AAV-mediated overexpression of *Kcnip4* in a humanized AD mouse model reduced the expression of the activity-dependent genes *Arc* and *c-Fos*, suggesting compensatory mechanisms against neuronal hyperexcitability. Our dataset provides a valuable resource for investigating mechanisms underlying resilience to neurodegeneration.

## MAIN

Advancements in single-cell omics have been pivotal in characterizing the transcriptomic diversity of the human neocortex and elucidating selective cell vulnerability in neurodegenerative dementias such as AD^1–6^. Single-nucleus profiling of the neocortex in AD has identified neuronal populations that are vulnerable and depleted early in the disease, such as layer 1 inhibitory interneurons expressing *NDNF/RELN* and layer 2/3 excitatory neurons expressing *CUX2/COL5A2*^2, 7, 8^. In contrast, few studies have focused on neuronal subtypes that, despite residing in similar microenvironments, remain preserved even in advanced stages of AD. Identifying these resilient subtypes and the mechanisms underlying their preservation could provide valuable insights for therapeutic strategies aimed at slowing disease progression.

We leveraged the progression of AD in the human neocortex—from association cortices to primary cortices^9–12^— to compare early-affected regions (prefrontal cortex, BA9; precuneus, BA7) with late-affected regions (primary visual cortex, BA17) using single-nucleus RNA sequencing (snRNA-seq). Although the neocortex follows a canonical 6-layer pattern, significant quantitative differences exist across regions^13–15^. For instance, layer 4 (L4) is expanded in primary sensory areas, while layers 2/3 and 5 (L2/3, L5) are more prominent in association cortices^3, 6, 16–19^. Comparing early- and late-affected areas thus provides a robust framework for examining cell-intrinsic and microenvironmental factors influencing selective vulnerability.

Neocortical L4, or the internal granular cell layer, is densely packed with small, granular neurons that serve as major postsynaptic targets of thalamic sensory nuclei and project locally or to nearby cortical regions. Its thickness varies considerably across different cortical areas, comprising 38% of the cortical ribbon in BA17 and 8.6% in BA9. In BA17, also known as the striate cortex, layer 4 contains a distinct band of myelinated fibers called the line of Gennari^17, 19^. L4 has long been considered a resilient area in AD due to its lower burden of tau in neurofibrillary tangles (NFTs), although it exhibits amyloid plaques^9, 20–22^. However, the composition of L4 at the single-cell level in AD progression remains poorly understood. In an unbiased manner, our study identified a resilient population of L4 neurons in the BA17 characterized by the co-expression of *RORB*, *CUX2*, and *EYA4*. Whether the resilience of these neurons is due to their specific connectivity, molecular properties, or interactions within the microenvironment remains unresolved, underscoring the importance of single-cell approaches in dissecting these complex factors and advancing research into neuronal resilience.

Our dataset includes samples from three neocortical regions (BA9, BA7, BA17) collected from 46 donors spanning all stages of disease progression (Braak stages 0–VI). To enrich for neurons, we performed fluorescence-activated nuclear sorting (FANS) for NeuN, resulting in 424,528 nuclei after quality control, of which 362,224 were neuronal. By integrating single-nucleus and spatial transcriptomics, we validated the layer-specific expression of 18 excitatory neuronal subtypes and identified the resilience of L4 neurons. We employed machine learning methods to validate neuronal subtype annotations across large-scale publicly available AD datasets^5, 8, 23^. Robust differential gene expression (DGE) analysis, utilizing linear mixed models, bootstrap resampling, and DESeq2 on pseudobulk aggregated counts, identified candidate genes associated with resilience. As proof of principle, we focused on *KCNIP4*, a gene encoding a voltage-gated potassium channel-interacting protein that regulates neuronal excitability in response to changes in intracellular calcium. We found that *KCNIP4* was upregulated in resilient L4 neurons during early disease stages. Furthermore, AAV-mediated delivery of *KCNIP4* in a humanized *App* knock-in AD mouse model (*App^SAA^*)^24^ reduced Arc and c-Fos expression, suggesting potential roles in regulating hyperexcitability. Our dataset is a valuable resource for investigating mechanisms of resilience in neurodegeneration.

## RESULTS

### Neuronal cell type composition during the spatiotemporal progression of AD in the neocortex

In the AD neocortex, neurodegeneration and tau pathology progress from association to primary cortices^9–12^. We profiled nuclear transcriptomes from 243 samples obtained from two association cortices (BA9, BA7) and one primary cortex (BA17) from 46 donors who died at various stages of disease progression and age-matched healthy controls (Braak stages 0–VI). The donors were categorized into three pathology groups—low, intermediate, and high (18, 10, and 18 donors, respectively)—based on neuropathological diagnoses using current consensus criteria^10^ (Fig. 1a; Supplementary Table 1). For each tissue sample, we collected two single-nucleus suspensions using FANS: one containing all nuclei and one enriched for neurons (NeuN^+^). In total, we profiled 655,407 nuclei. After rigorous QC to remove nuclei with low gene counts, high mitochondrial content, and doublets, we retained 424,528 high-quality nuclei for downstream analysis (Fig. 1b; Supplementary Table 1). The major cell types included 362,224 neurons (282,930 excitatory and 79,294 inhibitory), astrocytes (14,691), microglia (5,071), oligodendrocyte precursor cells (OPCs; 5,770), oligodendrocytes (36,589), and vascular cells (183) (Fig. 1c-f).

**Figure 1.**
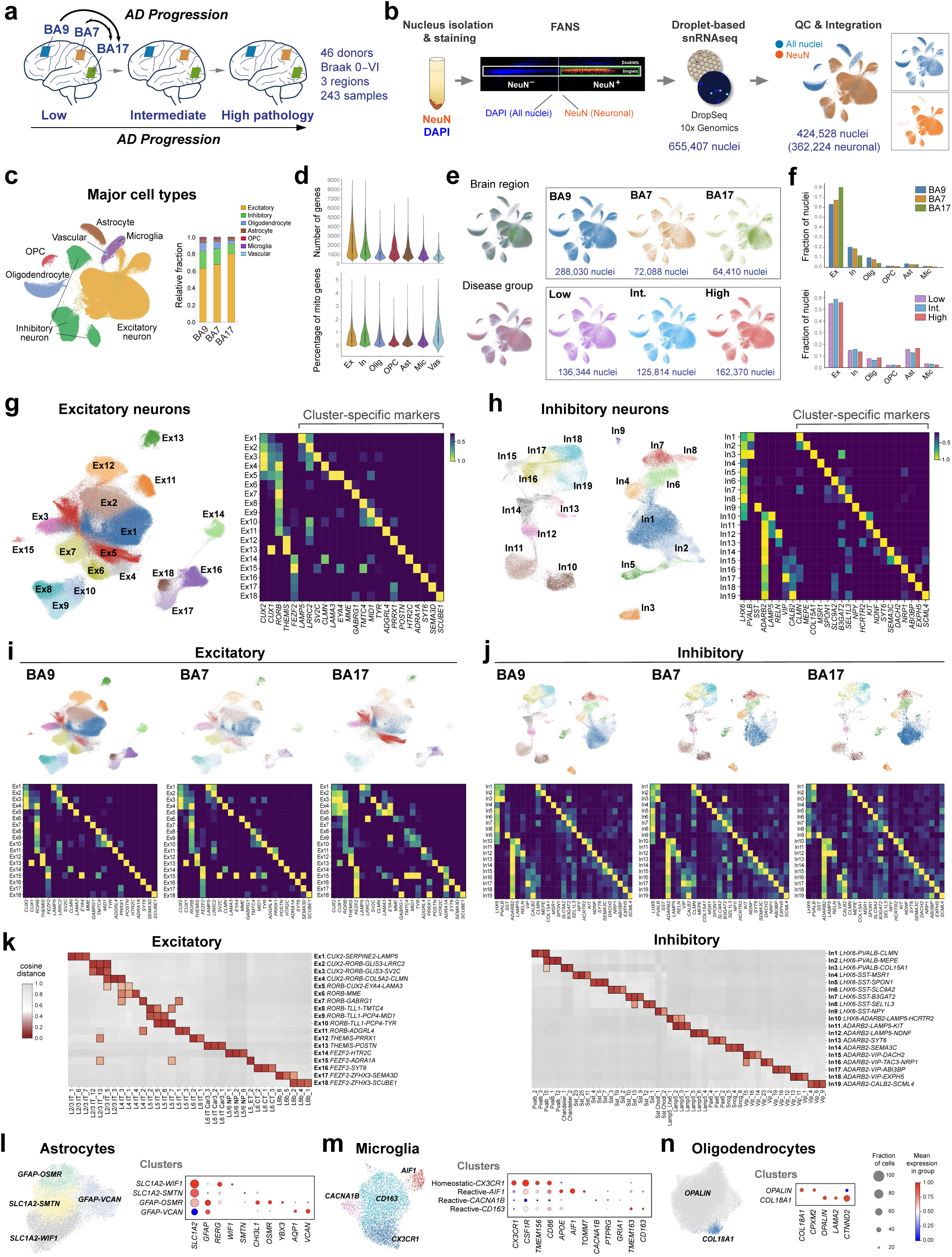
Neuronal cell composition across neocortical regions and AD pathology stages. **a**, Experimental design to study AD progression across neocortical regions and disease stages using snRNA-seq. **b**, Neuronal enrichment by FANS, snRNA-seq, and dataset integration yielded 424,528 nuclei (362,224 neurons, after QC). **c**, UMAP and bar plots representing the relative abundance of major cell types. **d**, Violin plots showing the median number of genes and percentage of mitochondrial genes within major cell types. **e**, UMAP plots splitting the datasets by region and disease stage group. **f**, Fraction of nuclei from each major cell type by region (top) and disease stage group (bottom). **g**,**h**, UMAP plots of the annotated excitatory and inhibitory clusters and heatmaps showing the normalized expression of selected subtype and cluster-specific marker genes. **i,j** UMAP plots and gene expression heatmaps for each brain region highlighting quantitative differences between association and primary cortices, and overall preserved marker genes across regions. **k**, Cosine distance matrix comparing the proximity in gene expression between the excitatory and inhibitory clusters from the SEA-AD DLPFC reference dataset^23^ (x-axis) and our BA9 dataset (y-axis). The closer the distance (lower values), the greater the similarity. The top three most closely related clusters are depicted. **l**−**n**, UMAP and dot plots showing the annotated glial subtypes and states.

We identified 18 excitatory (Ex) and 19 inhibitory (In) clusters, corresponding to neocortical neuronal subtypes, using stringent criteria. Our clustering strategy employed unsupervised Leiden clustering, combined with strict thresholds based on silhouette scores and Within Cluster Sum of Squares (WCSS), to enhance clustering reliability and ensure reproducibility. Clusters were named according to canonical markers for major subclasses (*CUX2*, *RORB*, *THEMIS*, and *FEZF2* for excitatory; *LHX6*, *ADARB2, PVALB*, *SST*, *VIP*, and *LAMP5* for inhibitory) along with 1−3 top marker genes for each cluster (Fig. 1g-h; Supplementary Table 2). Additionally, we selected gene sets (7−10 genes per subtype) whose combined expression precisely labeled each neuronal subtype across neocortical regions (Extended Data Figs. 1 and 2). The clusters and their marker genes showed consistent expression across BA9, BA7, and BA17. As expected, we observed significant differences in the abundance of neurons in specific excitatory clusters between association cortices and the primary visual cortex, reflecting their different cytoarchitecture^3, 13^. For instance, Ex5, characterized by the expression of *CUX2*, *RORB*, and *EYA4*, was overrepresented in BA17 (Fig. 1i). In contrast, all inhibitory clusters were well represented across the three regions (Fig. 1j).

We further assessed cluster reliability by comparing them with clusters from a large-scale AD reference dataset (Seattle Alzheimer’s Disease Brain Cell Atlas [SEA-AD])^23^, which includes nearly 1.4 million nuclei from the dorsolateral prefrontal cortex (DLPFC), using a cosine distance matrix to assess the similarity between the gene expression profiles of both datasets (Fig. 1k). Our annotations closely matched the reference dataset. Additionally, we used the semi-supervised single-cell ANnotation using Variational Inference (scANVI) model to annotate two AD reference datasets (SEA-AD DLPFC^23^ and Green and colleagues^5^), based on predictions from our 18 excitatory and 19 inhibitory neuron clusters. Our gene sets consistently labeled the clusters across datasets (Extended Data Figs. 1 and 2).

Glial cell states closely matched those from previous studies^2, 5, 25^ and included 4 astrocyte states, labeled by: *SLC1A2*/*WIF1* (homeostatic-protoplasmic), *GFAP/CHI3L1/OSMR* (reactive-inflammatory), *GFAP/AQP1/VCAN* (reactive-fibrous), and *SLC1A2/SMTN*; 4 microglia states: *CX3CR1* (homeostatic), *AIF1* or *CACNA1B* (reactive inflammatory), and *CD163* (reactive-phagocytic); and 2 oligodendrocyte states: *OPALIN* (myelinating) or *COL18A1* (immature) (Fig. 1k-m).

To spatially map the cortical layer distribution of the 18 excitatory clusters, we performed 10x Genomics Visium on 16 tissue sections from multiple neocortical regions in AD and control donors. We employed a custom-designed panel of 197 genes, which included cluster-specific marker genes identified from our snRNA-seq data, along with the commercial Human Neuroscience gene expression panel (1,186 genes) (Fig. 2a-c). We then integrated the spatial and single-nucleus transcriptomics data and applied variational autoencoders and deconvolution methods to quantify specific neuronal cell types based on the spatial distribution of marker genes. This approach allowed us to map the spatial distribution of each cluster on the tissue sections, with cortical layer boundaries delineated based on high-resolution H&E-stained images (Fig. 2a,b). Besides assigning layer identities to each cluster, this method highlighted regional differences, such as the overrepresentation of cluster Ex5 within L4 of BA17 (Fig. 2b). Thus, our integrated single-nucleus and spatial transcriptomics data identified robust clusters characterized by specific marker genes and gene sets, and mapped their spatial laminar distribution across neocortical brain regions (Fig. 2c,d).

**Figure 2.**
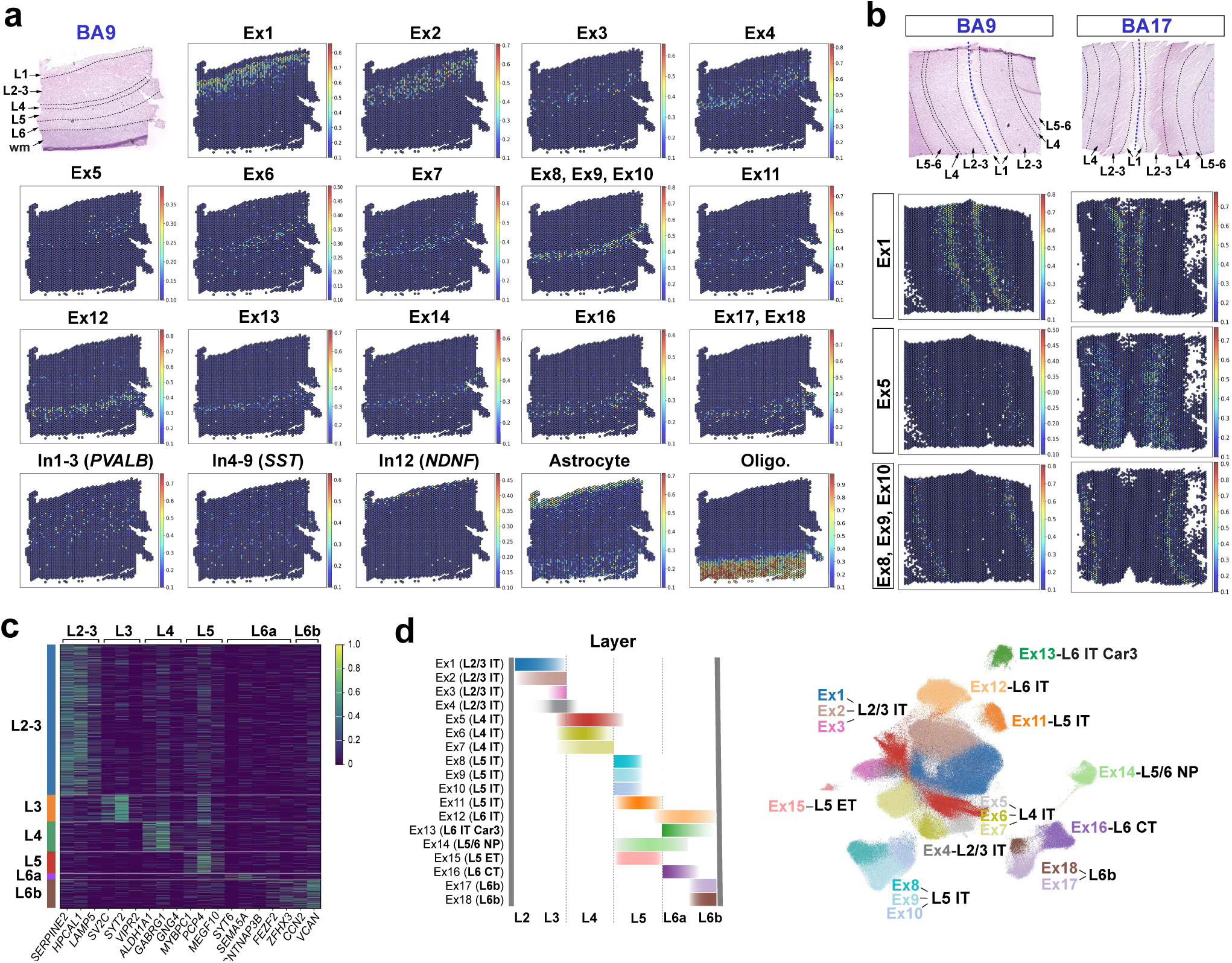
Layer-specific expression of excitatory neuronal subtypes. **a**, Spatial distribution of excitatory clusters in a representative BA9 section from a control donor. The H&E-stained tissue section indicates the boundaries between neocortical layers (top left). Expression of the excitatory subtypes annotated by snRNA-seq and visualized in the 10x Visium space using Stereoscope, highlighting the layer-restricted expression of each excitatory cluster. Smaller, overlapping clusters (Ex8/Ex9/Ex10 and Ex17/Ex18) are visualized together. *PVALB***^+^** interneurons span layers 2−5, *SST***^+^** interneurons span layers 1−6, whereas the *NDNF* interneuron subtype is restricted to L1. Astrocytes in L1 and in white matter and oligodendrocytes in white matter are shown for reference. **b**, Differences in L4 thickness and gene expression between BA9 and BA17. Representative H&E sections (top) and expression maps (bottom) illustrate the laminar expression of Ex5 (*RORB/CUX2/EYA4/LAMA3*) in L4, which is expanded in BA17, compared with Ex1 in superficial L2/3, and Ex8/Ex9/Ex10 in L5. **c**, Heatmap showing unsupervised layer distribution of top marker genes included in the 10x Visium gene panel. For example, *SERPINE2* and *LAMP5* label L2/3 cells, while *SV2C* and *SYT2* label deeper L3 cells. *ALDH1A1* and *GABRG1* label L4, and a subset of genes, including *PCP4,* label layers 3 and 5. Markers for L6 include *SYT6* and *SEMA5A* in L6a, and *ZFHX3*, *CCN2,* and *VCAN* in deep L6b. **d**, Summary of the 18 annotated excitatory subtypes and their layer-specific spatial distribution, incorporating annotations for excitatory neuron subclasses (L2/3 IT, L4 IT, L5 IT, L6 IT, L6 IT car3, L5/6 NP, L6b, and L6 CT) from the SEA-AD reference dataset^23^.

### Relative preservation of layer 4 excitatory neurons during AD progression

The thickness of neocortical L4 varies significantly across regions, comprising about one-third of the cortex in BA17^3, 17^. In advanced AD, L4 in BA17 shows amyloid plaques but minimal tau pathology^9, 20–22^. However, it remains unclear whether L4 is resilient across neocortical regions. To investigate this, we first identified marker genes for L4 excitatory neuronal subtypes (Ex5, Ex6, and Ex7; L4 IT). Consistent with previous studies, many genes expressed in L4 exhibited spatial gradients extending into layers 3 and 5^3, 16, 26, 27^. L4 was characterized by co-expression of *CUX2* (labeling L2−4) and *RORB* (labeling L3−5), with high expression of *CUX1*. The top cluster-specific markers for Ex5 included *EYA4, KCNH8*, *LAMA3, VAV3, KCNIP1,* and *TRPC3* (Fig. 3a). While Ex5 was particularly enriched in BA17, most of its marker genes were either absent or only detected in rare cells in BA9, except for *LAMA3*, which was expressed in a subset of Ex5 neurons across neocortical regions (Fig. 3b). Notably, all these Ex5 marker genes were also highly conserved in a reference dataset from the mouse neocortex^28^ (Fig. 3c). The other two clusters in L4, Ex6 (*RORB/MME*) and Ex7 (*RORB/GABRG1*), were both well represented in BA9 and BA7 but were rudimentary in BA17 (Fig. 3a,b, Extended data Fig. 1). While BA9-Ex5 did not share many marker genes with BA17-Ex5, Pearson correlation analysis of gene expression showed that BA9-Ex5 was more closely related to BA17-Ex5 than to other L4 clusters (Ex6 and Ex7) in BA9 or BA7 (Fig. 3d).

**Figure 3.**
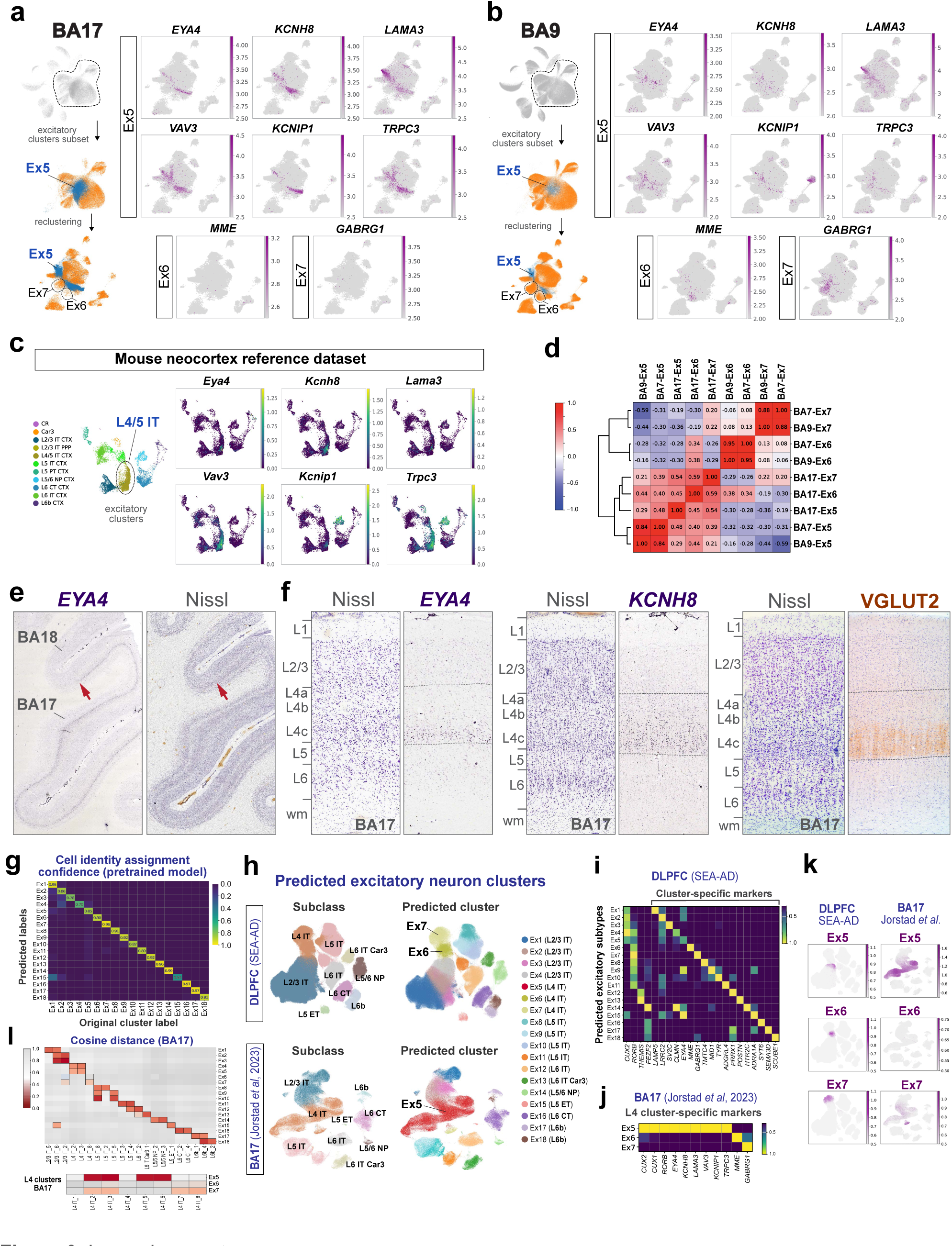
Markers of layer 4 across neocortical regions. **a,b,** UMAP plots highlighting the top L4 marker genes in BA17 (**a**) compared to BA9 (**b**). The Ex5 cluster (blue) and its top marker genes (*EYA4, KCNH8, LAMA3, VAV3, KCNIP1, TRPC3*) are overrepresented in BA17, whereas Ex6 (*MME*) and Ex7 (*GABRG1*) are overrepresented in BA9. **c**, UMAP plots from mouse neocortex snRNA-seq^28^ highlighting the conserved expression of top Ex5 marker genes in a cluster annotated as L4/5 intratelencephalic (IT). **d**, Pearson correlation coefficients measuring similarity in gene expression between L4 clusters (Ex5, Ex6, and Ex7) across different brain regions (BA9, BA7, and BA17). **e**,**f**, Identification of Ex5 neurons in L4 of BA17. Low-magnification images of the occipital cortex at the transition between BA17 and BA18 (primary and secondary visual cortex, respectively; red arrow in **e**) highlight the abundance of *EYA4*+ cells in BA17 (Allen Human Brain Atlas, https://human.brain-map.org/ish/experiment/show/80510718). Higher magnification images of BA17 show the expression of *EYA4* and *KCNH8* in L4 (Allen Human Brain Atlas, https://human.brain-map.org/ish/experiment/show/78937929). The boundaries of L4 are defined histologically in parallel Nissl-stained sections and by the expression of VGLUT2 in the terminals of thalamocortical projections from the LGN. **g**, Heatmap showing the assignment confidence scores for each excitatory cluster from the pretrained model using scANVI. **h**, UMAPs showing the author-annotated excitatory subclasses (left) and predicted excitatory clusters (right) in reference datasets from prefrontal cortex (SEA-AD DLPFC^23^) and primary visual cortex (Jorstad *et al*., 2023^3^). **i,j**, Gene expression heatmaps showing the normalized expression of cluster-specific marker genes in the predicted clusters from SEA-AD DLPFC (**i**) and BA17 L4 markers in the predicted clusters from subsetted L4 BA17 (**j**). **k**, Gene expression UMAPs of our cluster-defining gene sets for Ex5, Ex6, and Ex7 in the reference datasets from BA9 and BA17. **l**, Cosine distance matrix based on 3,000 highly variable genes from our BA17 dataset and the BA17 reference dataset^3^ demonstrating high similarity. In the top matrix, the three most closely related clusters, based on cosine distance, are depicted. In the bottom matrix, all L4 clusters are shown.

*In situ* hybridization (ISH) for *EYA4* and *KCNH8* in human BA17 tissue sections from the Allen Human Brain Atlas^29^ confirmed their expression in L4 and was consistent with labeling of L4 granule neurons. The highest expression was observed in deep layer 4c, with a sharp boundary at L5 (Fig. 3c,d), while expression in layers 4a and 4b was low. Notably, the most commonly used laminar nomenclature distinguishes three sublayers within L4: 4a, 4b, and 4c, although some authors classify layers 4b and 4c as part of L3, a view supported by tract-tracing studies in macaques^30, 31^. The expression of VGLUT2 (Fig. 3d), which labels presynaptic terminals from the lateral genicular nucleus (LGN) projecting to L4 in BA17 across species^32, 33^, matched the expression of *EYA4* and *KCNH8*. Therefore, *EYA4* and *KCNH8* preferentially label what is considered layer 4 proper in BA17.

To identify our L4 clusters in external datasets, we used scANVI to predict our annotations in three reference datasets from the prefrontal cortex (SEA-AD DLPFC^23^; Green *et al*., 2024^5^; Mathys *et al*., 2023^8^) and one from the primary visual cortex (Jorstad *et al*., 2023^3^) (Fig. 3e-j; Extended data Fig. 1b). We observed high similarity across datasets originating from the same brain region based on cosine distance scores, the expression of cluster-specific markers, and by plotting author-annotated and predicted clusters (Fig. 3g−l). As expected, the number of Ex5 cells predicted in the BA17 reference dataset was high: 63,870 cells (34.42%) out of 185,565 excitatory cells. In contrast, it was low in the prefrontal cortex reference datasets: 2,152 cells (0.33%) out of 660,751 excitatory cells in the SEA-AD dataset; 19,360 cells (3.03%) out of 637,968 in Green *et al*.; 3,361 cells (3%) out of 112,143 in Mathys *et al*., compared with 7,943 cells (4.36%) out of 182,140 in our BA9 dataset. Ex5 cells were most closely related to supertypes L4 IT_2, L4 IT_3, L4 IT_5, and L4 IT_6 from Jorstad *et al*., 2023^3^ (WithinArea_clusters) (Fig. 3l). In contrast, Ex6 (SEA-AD supertype L4 IT_4) and Ex7 (SEA-AD supertype L4 IT_2) were well represented in the prefrontal cortex across datasets and underrepresented in the BA17 dataset (Fig. 3h,k).

To evaluate neuronal loss in L4 relative to other cortical neuronal cell types, we analyzed neuronal composition in low, intermediate, and high pathology disease groups for each neocortical region (Fig. 4a,b Supplementary Table 3). The relative proportion of Ex5 L4 IT to total neurons increased significantly with disease progression, from ∼ 1.8% to 7.6% in BA9 and from ∼ 24% to 36.5% in BA17, comparing low to high pathology groups (Fig. 4a,c). The two other L4 excitatory populations, Ex6 L4 IT and Ex7 L4 IT, did not show significant changes. We obtained similar statistical results indicating the resilience of Ex5 using various methods, including a linear mixed model, Kruskal-Wallis test, beta regression, and scCODA—a model specifically designed to evaluate compositional changes in single-cell transcriptomic data^34^. Additionally, we observed a relative decrease in populations previously described as vulnerable, including L2/3 IT excitatory neurons (Ex1−Ex3), L5 IT excitatory neurons (Ex8), and a subpopulation of *SST* interneurons (In4) (Fig. 4a-c).

**Figure 4.**
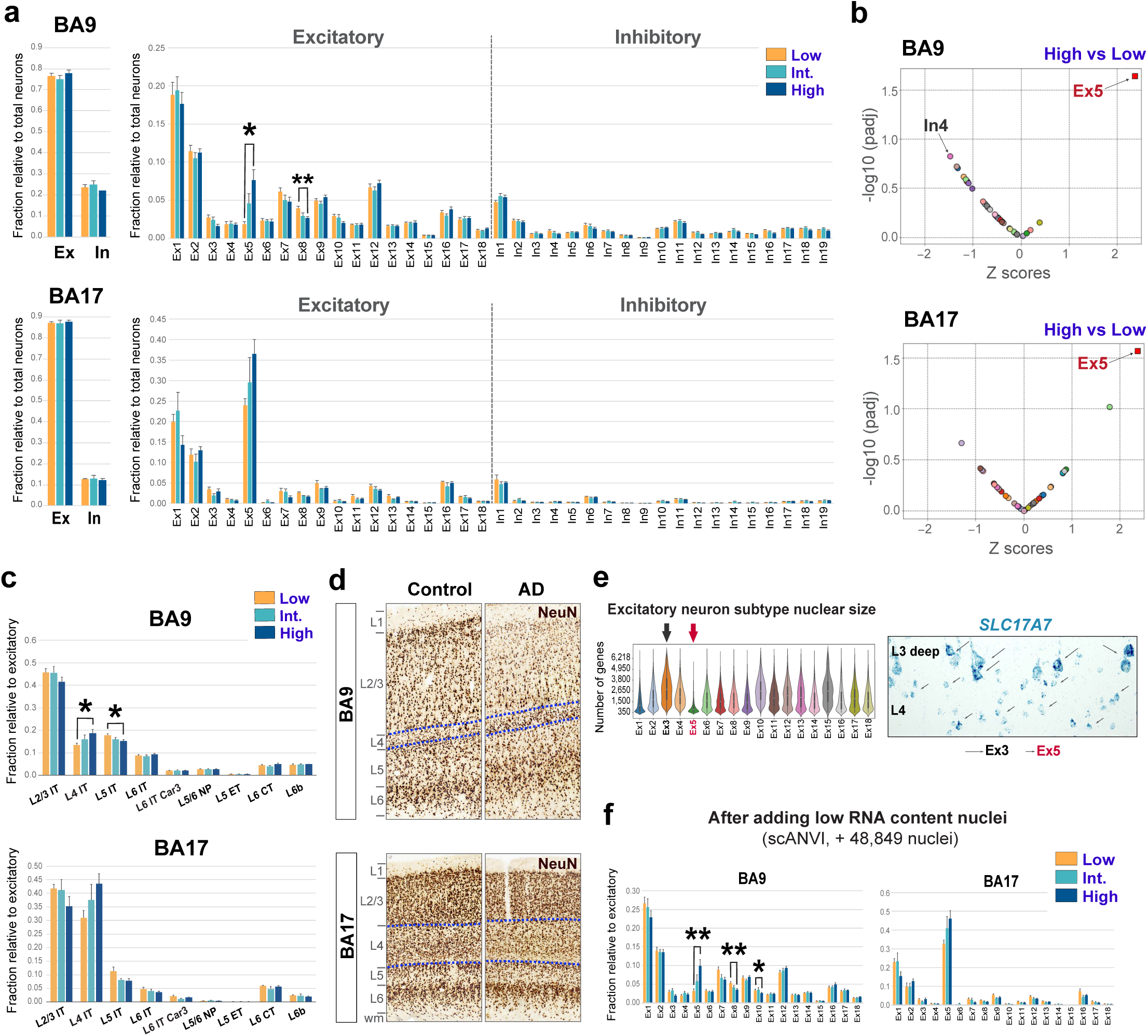
Relative preservation of layer 4 granular neurons in advanced AD. **a**, Fractions of total excitatory neurons and inhibitory neurons relative to all neurons (left plots) and of each excitatory and inhibitory neuronal subtype relative to all neurons (right plots), in BA9 and BA17, for each AD pathology group (low, intermediate, and high). The excitatory-to-inhibitory ratios were preserved, as previously reported^4^. Excitatory neurons in layers 2/3 and 5 tended to decrease with disease progression, while L4 neurons tended to increase. Changes were statistically significant for Ex5 (L4 IT; increased) and Ex8 (L5 IT; decreased) in the high compared with the low AD pathology group in BA9 (adjusted p-values = *p < 0.05, **p < 0.01; beta regression with correction for multiple testing). Bars represent mean proportions; error bars indicate standard error of the mean. **b**, Relative abundance of excitatory and inhibitory neuronal subtypes between low and high disease stages, quantified using a linear mixed model. Ex5 was significantly increased in both BA9 and BA17. In4 (SST^+^) was decreased but did not reach statistical significance in BA9 (adj p-value = 0.06). The x-axis represents z-scores; the y-axis represents significance (adj p-value). **c**, Cell proportions for each excitatory neuron supertype (L2/3 IT, L4 IT, L5 IT, L6 IT, L6 IT car3, L5/6 NP, L6b, and L6 CT) highlighting the relative depletion of L2/3 IT and L5 IT neurons and the preservation of L4 IT neurons in BA9 and BA17 (**\***p < 0.05; beta regression with correction for multiple testing). **d**, NeuN immunostaining on 50-μm-thick sections through the neocortex highlights the loss of pyramidal neurons in L2/3 and the relative preservation of granule cells in L4 in Braak VI AD compared with controls in BA9. Dashed lines delineate L4. **e**, Violin plots representing the number of genes within each cluster, which serve as an indicator of their relative neuronal size. In situ hybridization for *SLC17A7* in a tissue section from the BA9 illustrates the small size of layer 4 neurons (Ex5; red arrowheads) and the large size of deep layer 3 neurons (Ex3; black arrows). **f**, Annotation of previously discarded nuclei due to their low gene content (between 200 and 300 genes per nucleus) using scANVI. The heatmap shows prediction confidence for each excitatory cluster (x-axis: reference cells; y-axis: query cells). The bar plots show the fractions of each excitatory subtype relative to all excitatory neurons in BA9 and BA17 for each disease group. After incorporating 48,849 additional nuclei into the dataset, the relative preservation of Ex5 and depletion of L5 neuronal populations (Ex8, Ex10) in BA9 remained statistically significant (adjusted p-values = *p < 0.05, **p < 0.01; beta regression with correction for multiple testing).

Because Ex5 L4 IT neurons have small cell bodies and expressed up to 5-6 times fewer median number of genes compared to other excitatory neurons in the neocortex (Fig. 4d,e), we considered whether the QC filter for low gene counts (<300 genes) might have excluded them, potentially biasing our neuronal composition quantification. To address this, we selected previously filtered neuronal nuclei with gene counts ranging from 200 to 300 (62,498 nuclei) and used scANVI to predict their cell identities, using our high-quality dataset as a reference. After incorporating 48,849 nuclei (78%) that were confidently assigned (99% probability) to annotated clusters, we found that the overall neuronal composition remained unchanged, and the relative preservation of Ex5 neurons was statistically significant (Fig. 4f). Overall, our combined transcriptomic and histological data identified conserved marker genes for L4, revealing that the resilient Ex5 L4 IT population, which labels the internal granular cell layer, becomes increasingly prominent as neighboring neurons degenerate during disease progression. These findings were consistent across datasets and methods.

### Differential gene and pathway expression in vulnerable vs resilient neocortex in AD

To identify genes and pathways altered during disease progression in vulnerable and resilient regions, we performed DGE analysis comparing two disease stages (‘early’: low vs. intermediate pathology; ‘late’: intermediate vs. high pathology) and two neocortical regions (BA9 and BA17) for each neuronal subtype. Statistical power to detect differentially expressed (DE) genes is influenced by technical and biological factors, such as the number of nuclei, sequencing depth, RNA integrity, and age-dependent epigenetic changes^35, 36^. To address the heterogeneity of the samples and ensure the reliability of our findings, we applied several DGE methods, including a linear mixed model implemented in MAST and lme4, bootstrap resampling with 100 iterations, and DESeq2 on pseudobulk aggregated counts. (Fig. 5a, Extended data Fig. 3). We defined ‘high-confidence’ DE genes as those consistently identified across methods.

**Figure 5.**
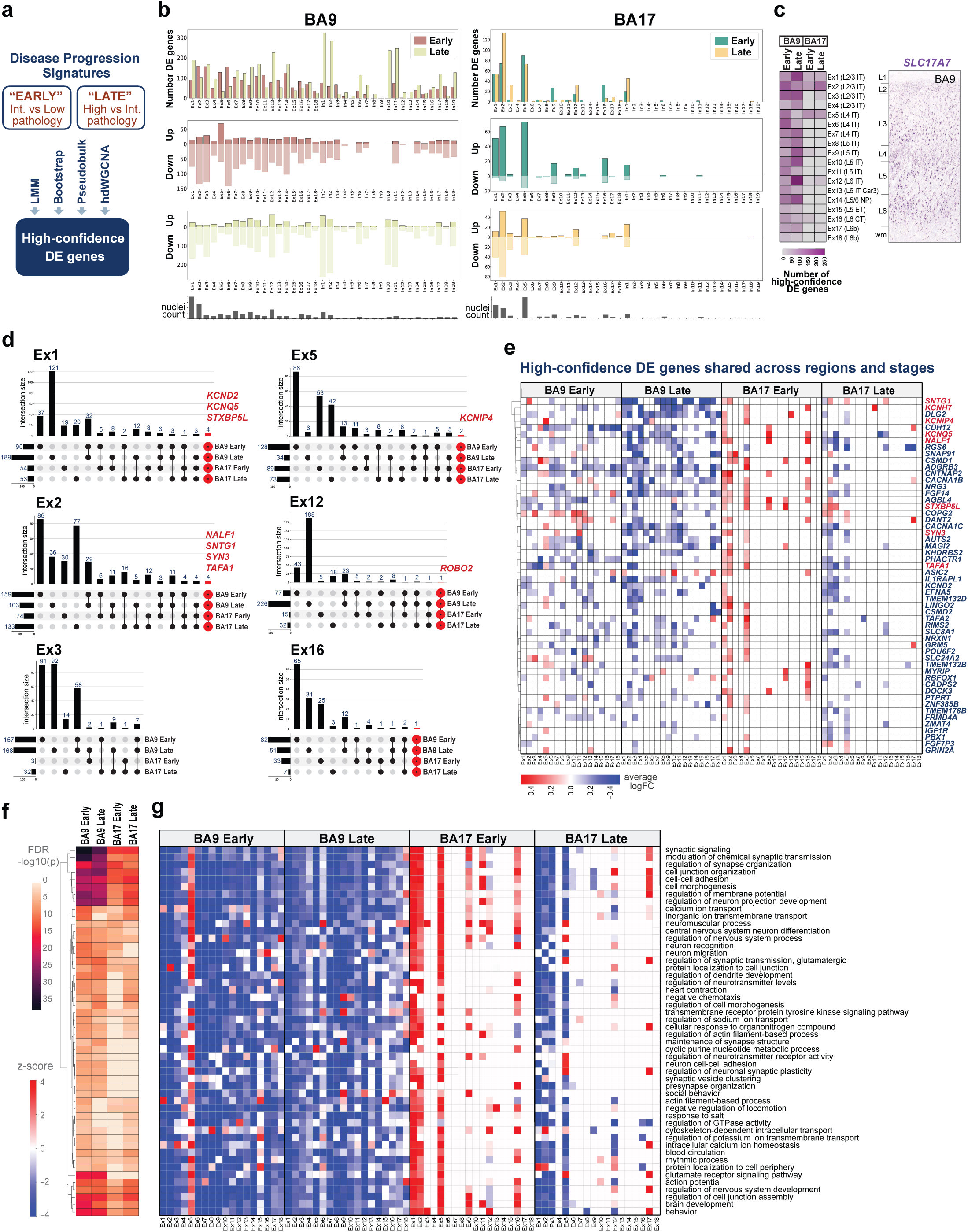
Transcriptome signatures of AD progression in neocortex. **a,** ‘High-confidence’ DE genes were identified using a linear mixed model and either bootstrap, pseudobulk, or hdWGCNA analyses. ‘Early’ and ‘late’ DE genes correspond to comparisons between intermediate vs low and high vs intermediate AD pathology groups, respectively. **b**, Bar plots illustrate the total numbers of DE genes, upregulated genes, and downregulated genes, identified by a linear mixed model within each excitatory and inhibitory neuronal cluster at early and late disease stages in the BA9 and BA17. Downregulation predominates; however, upregulation is relatively high at early stages in BA17. The number of nuclei per cluster is provided for reference. **c**, Heatmap depicting the number of high-confidence DE genes detected at high and late stages in BA9 and BA17 per excitatory cluster. The overall number of DE genes increases as pathology progresses from early to late stages and from BA9 to BA17. Notably, Ex5 exhibits a higher number of DE genes at early stages compared to late stages in both regions. The layer distribution of neuronal excitatory subtypes is shown in a section of BA9 stained with ISH for *SLC17A7* for reference. **d**, UpSet plots illustrate the intersection of high-confidence DE genes across BA9 and BA17 at early and late stages for six excitatory neuronal subtypes. Rows correspond to each of the four conditions, and columns represent the intersections. Bar charts depict the numbers of intersecting genes. Genes highlighted in red are differentially expressed in all four conditions. **e,** Gene expression heatmap of 54 high-confidence DE genes shared across brain regions and disease stages in excitatory neuronal subtypes. Only DE genes shared in at least 5 clusters are represented. Colors indicate the average log-fold change obtained from the linear mixed model. **f**,**g** Hierarchical heatmap visualization of functional enrichment analysis (**f**) in excitatory neurons from BA9 and BA17 at early and late stages highlights the common biological pathways enriched across regions and disease stages. High-confidence DE genes were used as input for gene ontology. The top 50 enriched pathways are represented. Heatmap visualization of the enriched pathways within each excitatory neuronal subtype (**g**) shows common patterns of gene downregulation (negative values; blue) in most subtypes from BA9 at both early and late stages and in BA17 L2-3 excitatory IT neurons (Ex1-3) at late stages, and gene upregulation (positive values; red) at early stages in BA17. Ex5 from both BA9 and BA17 at early stages share enriched pathways with upregulation in gene expression. The z-score values represent changes in gene expression.

The total number of DE genes was higher in BA9 compared to BA17 and in the ‘late’ disease groups compared to the ‘early’ groups, reflecting gene expression changes associated with AD progression (Fig. 5b,c, Supplementary Table 4). Subtypes previously identified as vulnerable, such as L2/3 IT excitatory neurons (Ex1, Ex2), exhibited more DE genes across both regions and disease stages. However, in BA9, some excitatory clusters, including the vulnerable Ex2 (L2/3 IT) and the resilient Ex5 (L4 IT), showed more significant changes in the ‘early’ compared to ‘late’ stages. Most DE genes in BA9 were downregulated, except for Ex5, where over 50% were upregulated in the ‘early’ stages. In contrast, in BA17, the majority of DE genes were upregulated, especially in the ‘early’ stages, in both vulnerable (Ex2) and resilient (Ex5) subtypes (Fig. 5b).

A total of 986 genes were categorized as ‘high-confidence’ DE genes. To distinguish between genes that were shared or unique across brain regions and disease stages (i.e., BA9-Early, BA9-Late, BA17-Early, BA17-Late), we generated UpSet plots showing intersections among these four conditions for each excitatory neuronal type (Fig. 5d). Although most genes were unique, likely due to the stringent criteria used to define ‘high-confidence’ DE genes, 15−27% were shared across at least two conditions within clusters with a high number of nuclei (Ex1, Ex2, Ex5, Ex12). The overlap of DE genes was greater between ‘early’ and ‘late’ disease stages within the same region than across brain regions, supporting the existence of vulnerability and resilient factors influenced by the microenvironment. Nevertheless, we identified 54 high-confidence DE genes common across all four conditions. Heatmaps of their expression changes revealed a consistent pattern: greater changes in BA9 compared to BA17, with downregulation increasing with disease progression in BA9 and upregulation shifting to downregulation with disease progression in BA17 (Fig. 5e). Genes exhibiting this pattern included *KCNH7*, *KCNQ5*, *DLG2*, *SNTG1, NALF1*, *CNTNAP2*, *FGF14*, *AUTS2*, and *MAGI2*. In contrast, a few genes, such as *COPG2* and *SLC24A2*, were upregulated at early stages in both BA17 and BA9. Notably, several high-confidence DE genes have previously been identified as genetic risk factors for AD, including *CSMD1, NRG3, SYN3, NRXN1, SLC24A2, DLG2,* and *KCNIP4*^37–42^.

Pathway enrichment analysis revealed shared pathways across regions and stages, including those involved in regulating synaptic organization, membrane potential, neurotransmitter levels, ion (calcium, sodium, and potassium) transport, intracellular calcium homeostasis, glutamate receptor signaling, synaptic vesicle clustering, and cell-cell adhesion (Fig. 5f,g; Supplementary Table 5). The same pattern persisted: enrichments were more significant in BA9 compared to BA17, and genes within the involved pathways were generally downregulated, except in the resilient regions (BA17-Early) and resilient neuronal subtypes (Ex5) (Fig. 5g).

### Genes and pathways associated with resilience in AD neocortex

To further define genes and pathways associated with resilience, we compared two neuronal subtypes: prototype vulnerable neurons (Ex2; L2/3 IT) and resilient neurons (Ex5; L4 IT) (Fig. 6a,b, Supplementary Table 6). We hypothesized that resilience-associated genes would be enriched and upregulated in Ex5, particularly at early stages and in BA17, consistent with disease progression and the preservation of L4 in AD. ‘High-confidence’ genes upregulated in Ex5 neurons at early stages in both BA9 and BA17 included: *CSMD1*, which encodes a synaptic protein that protects against complement-mediated synapse elimination^43^; *GRIN2A, GRM7, PTPRT, and KCNIP4,* which are involved in regulating neuronal excitability, synaptic transmission, synaptic organization, and synaptic plasticity; *SLC24A2*, a member of the calcium/cation antiporter superfamily involved in calcium homeostasis; *UBE2E2,* encoding an E2 ubiquitin-conjugating enzyme; *LINGO2*, a negative regulator of neuronal growth and survival; *TAFA1* and *TAFA2*, homologous genes encoding chemokine-like proteins with roles in neuronal survival; and *AUTS2*, involved in transcriptional activation and actin cytoskeleton reorganization. Some of these genes, such as *CSMD1, GRIN2A,* and *PTPRT,* were also upregulated in Ex2 and other excitatory neuronal subtypes at early stages in BA17, suggesting shared neuroprotective roles across different neuronal subtypes. Other DE genes upregulated early in BA17 and involved in synapse organization and function included: *CSMD2, NRXN1, NRG1, NRG3, TENM2, CACNA1B, GRID2, SLC8A1, SYN3, DLG2, DLGAP1, STXBP5L, NCAM2*, RIMS2, and *ADGRB3*. Additionally, genes upregulated at early stages in Ex5 included those encoding neurotrophic factors and proteins with neuroprotective properties, such as *NRG3, FGF14,* and *NCAM2* (Fig. 6a,b, Supplementary Table 6).

**Figure 6.**
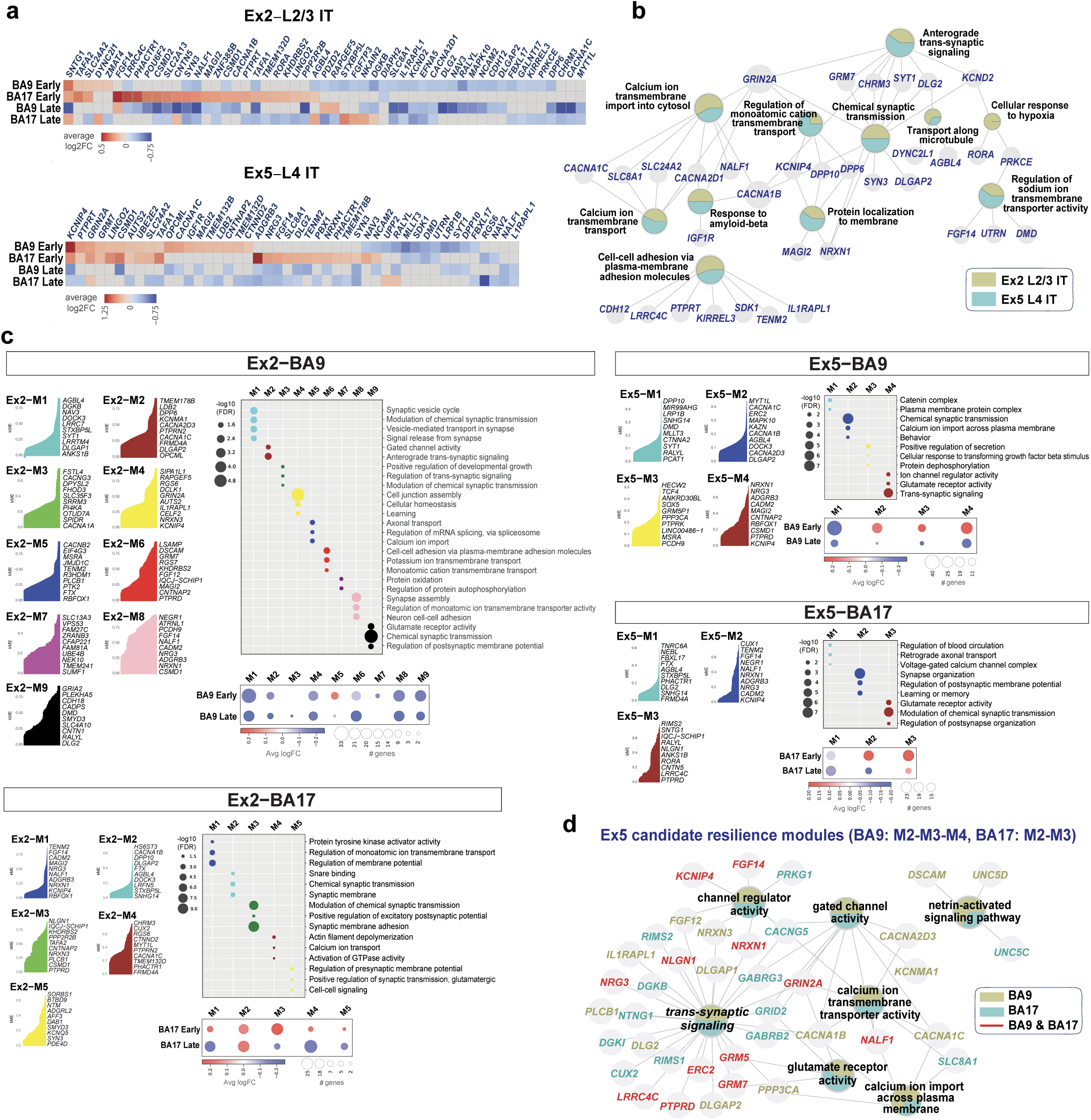
Transcriptome signatures of resilience in Ex5 L4 IT neurons. **a**, Heatmaps displaying ‘high-confidence’ DE genes shared across BA9 and BA17 at early and late stages in prototype vulnerable excitatory (Ex2; L2/3 IT) and prototype resilient (Ex5; L4 IT) neuronal subtypes. Genes differentially expressed in at least two of the four comparisons are depicted. Heatmaps are colored based on log2 fold change values. **b**, Biological function network of the genes represented in (**a)**. Colored nodes represent gene sets of biological functions contributed by the vulnerable (Ex2) and resilient (Ex5) subtypes. Node size reflects the number of connections between biological functions (minimum number = 5). **c**, Co-expression networks for vulnerable (Ex2; L2/3 IT) and resilient (Ex5; L4 IT) neuronal subtypes from BA9 and BA17, identified by hdWGCNA. The top 10 intra-module connected genes, ranked by Kme, for each module are represented. The enrichment dot plot illustrates the top functional categories of genes within each module. The color of the dots indicates the module, while the size of the dot reflects the significance of the enrichment. The gene expression dot plots represent the average logFC for each module at ‘early’ and ‘late’ disease stages. The size of the dot represents the number of differentially expressed genes, and the color indicates the magnitude of expression changes. **d**, Enrichment network for candidate resilient modules in Ex5 L4 IT neurons. The top 50 highly co-expressed genes from modules M2, M3, and M4 (BA9) and modules M2 and M3 (BA17), along with their enriched biological functions, are shown. Colors represent contributions from BA9 (moss), BA17 (teal), or both (red), along with their enriched biological functions.

Next, we analyzed high-dimensional weighted gene co-expression network analysis (hdWGCNA) data to compare systems-level changes in vulnerable (Ex2; L2/3 IT) and resilient neurons (Ex5; L4 IT) (Fig. 6c,d, Supplementary Table 7). In Ex5 neurons from BA17, we identified two candidate resilient modules, M2 and M3, where network genes were predominantly upregulated at early disease stages. The top 10 hub genes in these modules are: *KCNIP4, CADM2, NRG3, ADGRB3, NRXN1, NALF1, NEGR1, FGF14, TENM2,* and *CUX1* (for M2), and *PTPRD, LRRC4C, CNTN5, RORA, ANKS1B, NLGN1, RALYL, IQCJ−SCHIP1, SNTG1*, and *RIMS2* (for M3). For Ex5 neurons from BA9, we identified three candidate resilient modules: M2, M3, and M4 (Fig. 6c). A biological function network representation of these hdWGCNA genes, integrating the candidate resilience modules BA17–M2, M3 and BA9–M2, M3, M4, underscored the central roles of trans-synaptic signaling, calcium homeostasis, and neuronal excitability in resilience. Relevant genes within these modules include *GRIN2A, GRM5, GRM7, CACNA1B, CACNA1C, CACNG5, KCNIP4, NALF1, NRXN1, NLGN1, NRG3, PTPRD*, and *FGF14* (Fig. 6d).

### Increased *KCNIP4* expression is associated with resilience in AD

We focused on *KCNIP4*, a gene specifically upregulated in resilient Ex5 neurons at early disease stages in both BA17 and BA9 (Fig. 6a), as a proof of principle to validate our approach for identifying genes associated with resilience. This gene encodes a voltage-gated potassium channel-interacting protein (KCHIP4 or KCNIP4) that regulates neuronal excitability. KCNIP4 also interacts with Presenilins and has been previously linked to AD^44, 45^. Our analysis showed that *KCNIP4* is predominately expressed in excitatory neurons (except Ex14; L5/6 NP) and OPCs (Fig.7a), as well as a microglia cluster characterized by high expression of synapse-related genes (cluster Microglia-Reactive-*CACNA1B;* Supplementary Table 2). Using ANCOVA (Analysis of Covariance) to estimate *KCNIP4* expression across disease stage groups while controlling for fixed covariates (assay, sex, RIN, total counts) and random effects (donor), we consistently observed increased *KCNIP4* expression in Ex5 as disease progressed (Fig. 7b).

**Figure 7.**
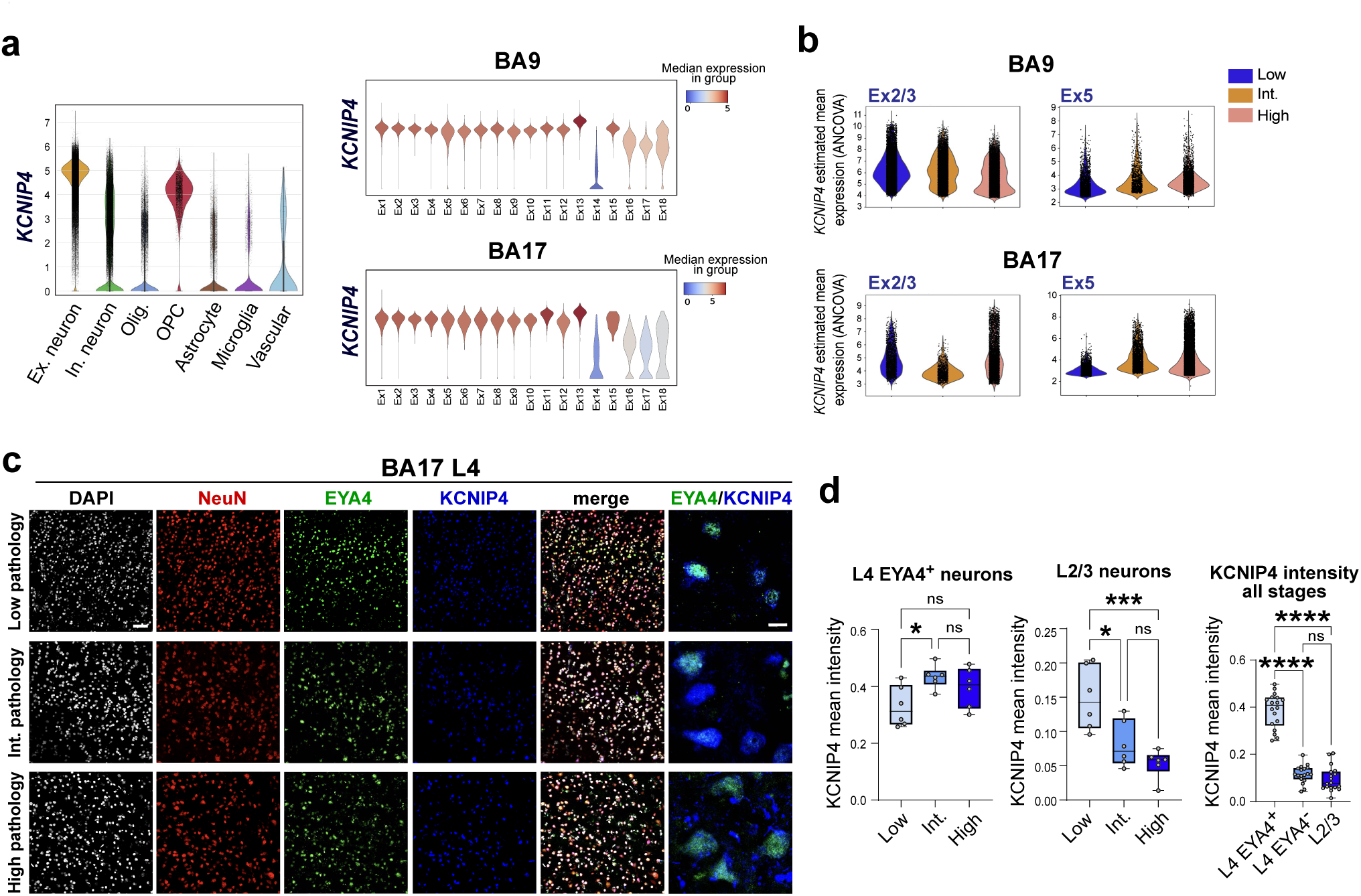
KCNIP4 upregulation in resilient L4 neurons. **a**, Violin plots showing *KCNIP4* gene expression across major cell types (left) and excitatory neuronal subtypes from BA9 and BA17 (right). **d**, Violin plots showing *KCNIP4* gene expression levels across disease stages (low, intermediate, and high pathology) in Ex2/3 and Ex5 neurons from BA9 and BA17. Estimated mean gene expression values were derived from a linear mixed model incorporating analysis of covariance (ANCOVA). **c**, Immunostaining for KCNIP4, EYA4, and NeuN in cryosections from low, intermediate, and high pathology stages illustrating increased expression of KCNIP4 in L4 EYA4^+^ neurons in BA17. **d**, Quantification of KCNIP4 protein expression levels in L4 EYA4^+^ neurons, L4 EYA4**^−^** neurons, and L2/3 neurons from BA17 across disease stages (n = 6 donors per disease group). Data are shown as median ± IQR. ANOVA with Tukey’s test was used for multiple comparisons (*p < 0.05; ***p < 0.001; ****p < 0.0001). Scale bar = 30 µm.

To quantify KCNIP4 protein levels in resilient versus vulnerable neurons, we performed immunohistochemistry for KCNIP4, EYA4, and NeuN in sections of BA17 from low, intermediate, and high pathology groups (Fig. 7c). EYA4 labels L4 granule cells in the cerebral cortex and is also expressed by a subset of GABAergic interneurons, which are sparse and located predominantly in the superficial layers. The mean intensity of KCNIP4 in neuronal somas was significantly higher in L4 EYA4^+^ neurons at intermediate disease stages compared to controls, and lower in supragranular (L2/3) neurons at intermediate and high stages compared to controls (Fig. 7d).

To evaluate *KCNIP4* upregulation in excitatory neurons, we induced *Kcnip4* expression using AAV-mediated delivery in a humanized *App* knock-in mouse model of familial AD (*App^SAA^* KI/KI)^24^. We generated the AAV vector PHP.eB-CaMKIIa-Kcnip4-P2A-EGFP, using the PHP.eB serotype to efficiently transduce neurons in the CNS, the CaMKIIa promoter to selectively target excitatory neurons, the mouse *Kcnip4* transcript, and the P2A-EGFP sequence as a reporter. As a control, we used the same AAV containing only EGFP (Fig. 8a). To evaluate the ability of the *Kcnip4* AAV to increase KCNIP4 protein levels in the mouse brain, we performed Western blotting on cortex tissue lysates from 12-month-old WT mice treated with 2 different doses of *Kcnip4* AAV (5 × 10^10 vg and 1 × 10^11 vg, retroorbitally). Mice treated with the higher dose showed a significant increase in KCNIP4 (Fig. 8b). We injected 12-month-old homozygous *App^SAA^*mice, which exhibit amyloid plaques, microgliosis, and plaque-associated dystrophic neurites^24^, with *Kcnip4* AAV and control AVV (1 × 10^11 vg, retroorbitally). WT mice from the same genetic background and age received control AAV. Mice were sacrificed, and brain tissue was collected one month after injection. GFP^+^ neurons were detected throughout the cerebral cortex, and to a lesser extent in the hippocampus (Fig. 8c). To estimate transduction efficiency, we quantified the percentage of GFP^+^ neurons in cerebral cortex. In the three animal groups, GFP labeled approximately 10% of total neuronal population (Fig. 8d). AD pathology in the treated mice was not significantly modified by *Kcnip4* overexpression, as no significant differences were found in amyloid plaques (determined by an anti-human amyloid beta antibody, Fig. 8e) or in reactive astrogliosis (determined by GFAP, Fig. 8f). A trend toward decreased microgliosis was observed (as determined by IBA1 staining), although it did not reach statistical significance (p = 0.08) (Fig. 8g).

**Figure 8.**
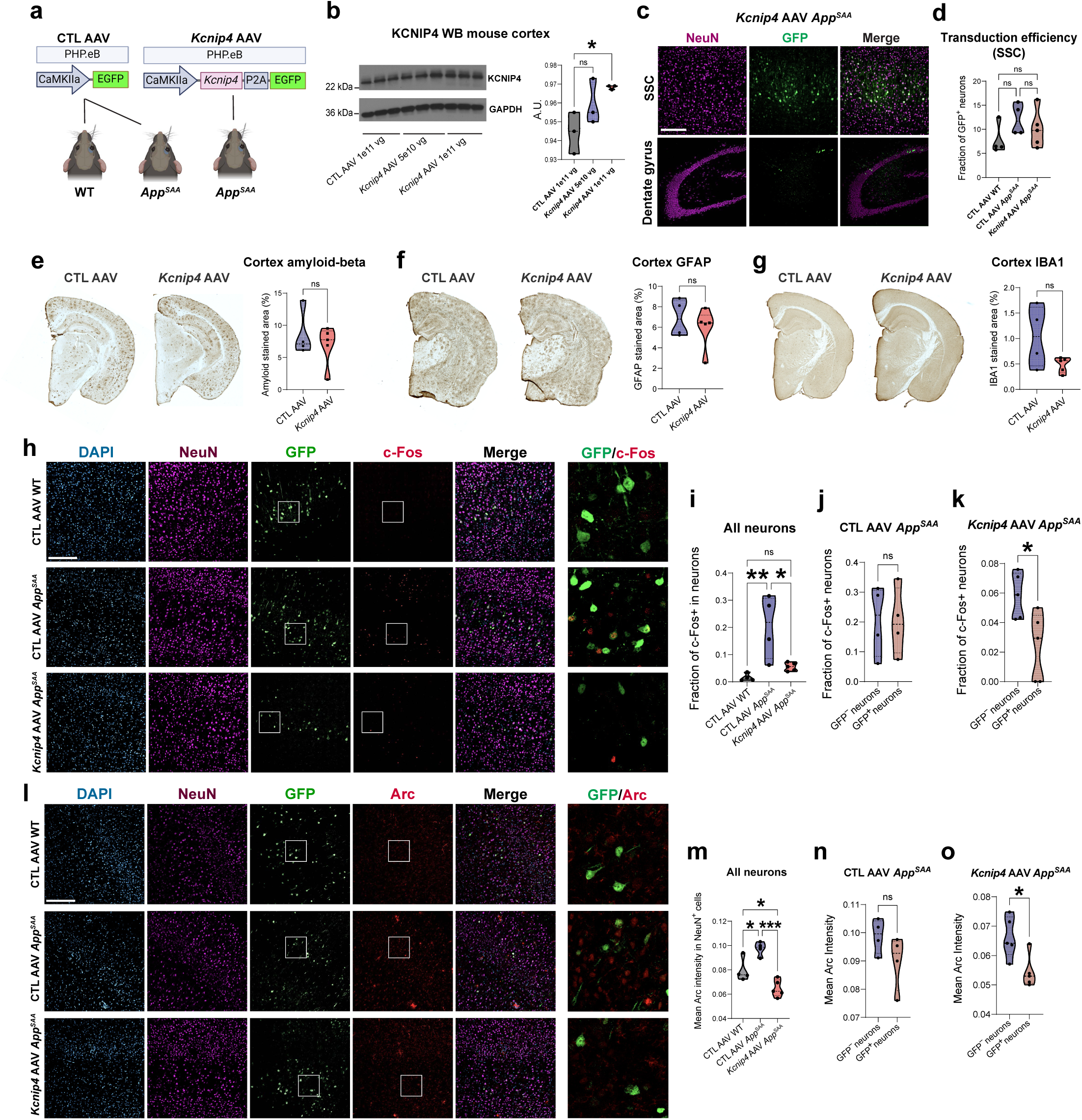
AAV-mediated delivery of *Kcnip4* in excitatory neurons reverses markers of hyperexcitability in a humanized mouse model of AD. **a**, Graphical representation of the AAVs and mice used in the study. All mice were 12 months old at the time of AAV retroorbital injection and were sacrificed one month later. **b**, Western blot quantification and representative image of KCNIP4 levels in cerebral cortex lysates following two different doses of *Kcnip4* AAV (n = 3 per group). **c,d**, Representative images of cerebral cortex and hippocampus from *Kcnip4* AAV-treated mice and quantification of transduction efficiency of the different AAVs in somatosensory cortex (SSC) in WT and *App^SAA^* mice. **e−g**, Representative images and quantification of cortical amyloid beta, GFAP, and IBA1 immunostaining in *App^SAA^* mice treated with *Kcnip4* AAV or control AAV. **h−k**, Quantification of c-Fos in *App^SAA^* mice treated with *Kcnip4* AAV or control AAV, and WT mice treated with control AAV: **h**) Representative immunofluorescence images through the SSC co-stained with DAPI, NeuN, GFP, and c-Fos; **i**) Fraction of c-Fos-positive cells in all cortical neurons across study groups; **j,k**) Fraction of c-Fos-positive cells in GFP^+^ compared to GFP^-^ neurons. **l−o**, Quantification of Arc immunostaining in *App^SAA^* mice treated with *Kcnip4* AAV or control AAV, and WT mice treated with control AAV: **l**) Representative immunofluorescence images through the SSC co-stained with DAPI, NeuN, GFP, and Arc; **m**) Mean Arc staining intensity in all cortical neurons across study groups; **n,o**) Mean Arc staining intensity in GFP^+^ compared to GFP^-^ neurons. All data are shown as median ± IQR (n = 5 for *Kcnip4* AAV in *App^SAA^*mice, and n = 4 for control AAV in WT or *App^SAA^* mice). A t-test was used for pairwise comparisons, and ANOVA with Tukey’s test was used for multiple comparisons (*p < 0.05; **p < 0.01, ***p < 0.001). Scale bars = 200 µm.

KCNIP4 is an integral component of Kv4 channel complexes and belongs to the EF-hand family of small calcium-binding proteins. Like other Kv channel-interacting proteins, it may control neuronal excitability by regulating A-type outward potassium currents^46^. Thus, we hypothesized that increased KCNIP4 expression may reduce neuronal hyperexcitability in AD. To investigate this, we quantified c-Fos and Arc, two immediate-early genes widely used as markers of neuronal activation^47^. These markers increase with excessive neuronal stimulation and seizures and have been shown to be altered in AD^48^. Using GFP as a marker of transduced neurons, we found that *Kcnip4* AAV-mediated delivery in 12-month-old *App^SAA^* mice reduced the proportion of c-Fos^+^ neurons in the GFP^+^ compared to GFP^-^ populations (Fig. 8h-k). No changes in c-Fos proportions were observed in *App^SAA^* mice treated with control AAV. When comparing all cortical neurons in *App^SAA^*and WT mice, we found that *App^SAA^* exhibited increased levels of c-Fos, which were reversed by *Kcnip4* AAV (Fig. 8i-k). We observed similar results for Arc expression, with reduced intensity in GFP^+^ compared to GFP^-^ neurons in *App^SAA^* mice treated with *Kcnip4* AAV, and a reversal in Arc expression in treated *App^SAA^*mice compared to WT controls (Fig. 8l-o). Thus, increased KCNIP4 expression in excitatory cortical neurons in a humanized mouse model of AD reduced c-Fos and Arc, markers of neuronal activation and hyperexcitability, suggesting a role in promoting resilience against hyperexcitability in AD.

## DISCUSSION

Our strategy leveraged the spatiotemporal progression of AD to explore cellular resilience. The primary visual cortex exhibits mild degeneration, even in end-stage AD, yet it has not been fully explored for potential resilience factors^9–11, 49^. Layer 4 neurons, considered resilient in AD due to low tau pathology, have not been consistently characterized in previous snRNA-seq studies. We identified a cluster, Ex5, enriched in BA17 and comprising granular L4 IT neurons, that remains relatively preserved in early-and late-stage AD cortices. This resilience was linked to upregulated genes related to synaptic function and calcium homeostasis, including *KCNIP4*, suggesting compensatory mechanisms against hyperexcitability—an early feature in AD pathogenesis observed in human and animal models^50–52^.

Layer 4 cytoarchitecture varies across neocortical regions and is highly specialized in areas receiving topographic sensory input from the thalamus, such as BA17^19, 53^. BA17 is relatively thin, highly myelinated, and has a low interneuron-to-excitatory neuron ratio. It features an expanded L4 with high neuronal density and distinct sublayers that receive input from the LGN and project to nearby regions. Morphologically, BA17 contains different types of excitatory neurons: a dominant population of granular neurons enriched in L4c, a smaller population of pyramidal neurons enriched in L4a and L4b, and rare “giant” stellate cells arranged horizontally in L4b. In association neocortex, L4 is thin but visible, appearing discontinuous and segmented into vertical columns in some regions, blending with pyramidal neurons of layers 3 and 5^17, 31^. Our analysis identified three distinct molecular subtypes across neocortical regions: Ex5 (*CUX2/RORB/EYA4/LAMA3*), Ex6 (*RORB/MME*), and Ex7 (*RORB/GABRG1*), which we validated across publicly available datasets^3, 5, 8, 23^. Ex5 was overrepresented in BA17, while Ex6 and Ex7 were more prominent in association cortices (BA9, BA7). We found that the Ex5 cluster-defining genes *EYA4* and *KCNH8* preferentially label granule neurons in deep layer 4c, the same area receiving VGLUT2+ terminals from the LGN. Previous snRNA-seq studies of BA17 from healthy individuals have identified specialized L4 excitatory neuron subtypes with greater granularity. The Ex5 cluster closely matches L4_IT3, a dense pan-L4 marker, and includes L4_IT2 and L4_IT5, enriched in layers 4cβ and 4cα, respectively^3^. Our analysis validates L4 excitatory neuronal subtypes across neocortical regions and stages of AD progression, providing a framework for identifying gene expression changes associated with resilience.

In excitatory neurons, gene upregulation under hyperexcitability-associated conditions, such as AD, may indicate pathological hyperexcitability, a maladaptive cellular response, or a compensatory response aimed at preventing excitotoxic damage. For instance, snRNA-seq profiling of cortical biopsies from living subjects with early pathology revealed molecular and electrophysiological properties indicative of hyperactivity in L2/3 pyramidal neurons prior to their loss. This study identified the upregulation of *APP, PRNP, ATP1A3, SNAP25, SYT1,* and *CDK5* as a signature of pathological hyperexcitability in vulnerable L2/3 IT neurons. In contrast, in resilient L4 IT neurons, we observed the upregulation of genes associated with neuroprotection—*GRIN2A, RORA, NRXN1, NLGN1, NCAM2, FGF14, NRG3, NEGR1,* and *CSMD1*—suggesting resilience and broadening our understanding of gene expression changes associated with hyperexcitability in AD.

Our analysis revealed an early upregulation of *KCNIP4* in resilient Ex5 L4 IT neurons; in contrast, *KCNIP4* was downregulated in vulnerable Ex2 L2/3 IT neurons during stages of cell death, with an overall decline observed in late-stage disease. *KCNIP4* is a member of the K-channel interacting proteins (KChIPs), which include KChIP1, KChIP2, KChIP3 (DREAM/calsenilin), and KChIP4 (CALP), encoded by the *KCNIP1-4* genes. KChIP4 interacts with Kv4.2 channels, a subtype of voltage-gated potassium channels that are key regulators of neuronal excitability. KChIP4 expression influences the subcellular localization and biophysical properties of Kv4 channels. Increased binding of KChIP4 enhances the recovery from inactivation of Kv4.2, thereby exerting an inhibitory effect on neuronal excitability ^54^. Additionally, KCNIP4 interacts with presenilins PSEN2 and PSEN1, potentially modulating APP processing and Aβ levels ^44, 55^. Notably, KCNIP4 belongs to the recoverin branch of the EF-hand superfamily, characterized by four EF-hand calcium-binding motifs. Several members of this family have demonstrated neuroprotective properties^56^. Thus, KCNIP4’s role in regulating neuronal excitability and APP processing may confer neuroprotection against excitotoxicity within hyperexcitable networks, particularly in response to elevated intracellular calcium levels.

Hyperexcitability has also been implicated as a pathogenic mechanism in other neurodegenerative diseases, such as amyotrophic lateral sclerosis and Huntington’s disease, and is associated with aging. For example, hyperexcitability in sleep circuits can lead to sleep instability and fragmentation, particularly in older adults ^52, 57–59^. Thus, hyperexcitability may serve as an early biomarker of neurodegeneration and a therapeutic target. Recent interventions targeting neuronal hyperexcitability in AD include the antiepileptic drug levetiracetam and emerging non-pharmacological brain stimulation techniques^60–62^. Our study identifying neurons preserved in end-stage AD and genes associated with neuronal excitability in these cells, such as *KCNIP4*, provides insights into cellular resilience in neurodegeneration and may guide the development of interventions to slow disease progression.

## METHODS

### Postmortem brain tissue

Brain tissue from AD and age-matched control donors was obtained from the UCLA Department of Pathology and Easton Center, the NIH Neurobiobank (Sepulveda repository in Los Angeles, CA, and Mt. Sinai Brain Bank in New York City, New York), and Stanford’s Department of Pathology and Alzheimer’s Disease Research Center. AD neuropathology was evaluated by a neuropathologist using the ABC score (National Institute on Aging and Alzheimer’s Association Research Framework criteria)^10^. Relevant information such as age, sex, ethnicity, brain weight, and postmortem interval (PMI) were recorded when available. *APOE* genotyping was performed using the SNP Genotyping service from Genewiz (Azenta Life Sciences) with genomic DNA isolated from fresh-frozen brain tissue samples. No cases with imaging or gross findings consistent with large vessel territorial infarction, hemorrhage, primary or metastatic neoplasms, or CNS infection were included. Cases with histological evidence of hypoxic-ischemic brain injury were excluded. Tissue blocks selected for snRNA-seq underwent immunohistochemical assessment, including H&E and Nissl stains to confirm tissue integrity and the absence of microinfarcts or other focal pathologies. NeuN immunohistochemistry was performed to confirm the absence of decreased NeuN immunostaining, which could bias the sorting of NeuN^+^ neuronal nuclei by FANS. Tau and amyloid immunohistochemistry were also performed to assess the extent of pathology in the same blocks utilized for snRNA-seq.

The tissue samples were collected from three regions: the prefrontal cortex (BA9), precuneus (BA7), and primary visual cortex (BA17), encompassing all stages of disease progression. A total of 46 donors contributed to the study (42 for BA9, 15 for BA7, and 24 for BA17). The stages of disease progression were categorized into three groups: low pathology (18 donors; 6 females, 12 males), intermediate pathology (10 donors; 7 females, 3 males), and high pathology (18 donors; 12 females, 6 males). The criteria for each group were based on the presence and distribution of tau aggregates, according to the Braak staging system^9^, and of amyloid pathology, including diffuse and neuritic amyloid plaques. The density of neuritic amyloid plaques was semi-quantified using the CERAD (C) staging system^63^. The low pathology group included cases with no tau or amyloid pathology, with low AD neuropathologic change (AD), and cases of primary age-related tauopathy (PART), a pathology associated with aging that features NFTs with similar morphology and distribution as in AD in the absence of amyloid^64^. The PART cases in this study had a Braak stage I−III. The intermediate pathology group included cases with Braak stage III−IV and diffuse plaques or sparse (C1) neuritic plaques. The high pathology group included cases with Braak stage V−VI and moderate (C2) or abundant (C3) neuritic plaques. The mean age of the donors in the low, intermediate, and high pathology groups were 70.5 ± 9.2, 81.9 ± 13.6, and 82.4 ± 10.4 years, respectively.

RNA integrity number (RIN) was measured in all the tissue blocks selected for snRNA-seq. Total RNA extraction from ∼20 mg of tissue was performed using Trizol reagent followed the RNeasy Plus Mini kit (Qiagen cat # 74134) according to the manufacturer instructions. Purified RNA was quantified using the Agilent Bioanalyzer 2100 RNA Nano chips (Agilent Technologies cat # 5067-1511) according to the manufacturer instructions. There were no significant differences in the RIN (5.8 ± 0.7, 6.2 ± 0.7, and 6.2 ± 0.7, respectively) and in the PMI (15.6 ± 8.2, 12.8 ± 8.2, and 13.8 ± 9.7 hours, respectively) between the low, intermediate, and high pathology groups.

### Single nuclear isolation and neuronal nuclei enrichment

The fresh-frozen brain tissue blocks (∼3 ξ 2 ξ 0.5 cm) were stored at −80°C. Adequate orientation of the blocks was ensured to enable full-thickness sectioning of the cortical ribbon with a proper representation of all layers. To that end, thick sections (∼500 µm) were cut spanning the entire thickness of the cerebral cortex, from the leptomeninges to the underlying white matter. The cryostat was set at −12°C to facilitate the cutting of these thick sections while preserving the remaining tissue block frozen for further experiments. Under a stereomicroscope, the tissue slices were dissected to remove the white matter and leptomeninges. For each experiment, ∼100 mg of cortical gray matter was utilized. To prevent further RNA degradation, all subsequent steps were conducted on ice under RNase-free conditions. The tissue was chopped into small pieces (< 1 mm^3^) using a chilled razor blade and homogenized with a Dounce tissue grinder (Kimble cat # 885300-0007). Each tissue sample was dissociated in 2.4 mL of homogenization buffer containing 10 mM Tris pH 8, 5 mM MgCl_2_, 25 mM KCl, 250 mM sucrose, 1 μM DTT, 0.5x protease inhibitor (cOmplete, Roche cat # 46931590010), 0.2 U/μL RNase inhibitor, and 0.1% Triton X-100. Typically, ∼30 grinder strokes with pestle B (0.020−0.056 mm clearance) were required. Microscopic examination using a hemocytometer was conducted to assess the number of nuclei and the presence of clumps and debris. The homogenates were subsequently filtered through a 40-μm cell strainer and transferred into two 1.5-mL Eppendorf tubes.

Iodixanol gradient centrifugation was used to further clean-up the nuclei and remove myelin debris. The homogenate was first centrifuged at 1000 ×g for 8 min at 4°C. The supernatant was discarded, and the pellets were gently resuspended in 450 μL of homogenization buffer. An equal volume (450 μL) of 50% v/v iodixanol medium (41.25 mM sucrose, 24.75 mM KCl, 5 mM MgCl2, 10 mM Tris [pH 8], and 50% v/v iodixanol) was added to the homogenate and gently mixed with a pipette. The mixture was then transferred to a new 2-mL Eppendorf tube containing 900 μL of 29% iodixanol medium (125 mM sucrose, 75 mM KCl, 15 mM MgCl2, 30 mM Tris [pH 8], and 29% v/v iodixanol) by slow layering on the top. The tubes were centrifuged at 13,500 ×g for 20 min at 4°C, resulting in the sedimentation of nuclei. The top layer, containing abundant myelin, and the supernatant were removed and discarded carefully, avoiding contamination of the nuclei pellet. The pellets were detached by carefully pipetting with ∼50 μL of immunostaining buffer (0.1 M phosphate-buffered saline [PBS; pH 7.4], 0.5% bovine serum albumin [BSA], 5 mM MgCl_2_, 2 U/mL DNAse I, and 0.2 U/μL RNase inhibitor), transferred to clean tubes, and gently resuspended in a total volume of 200 μL of immunostaining buffer. After a 15-min incubation with immunostaining buffer, at 4°C, with gentle rocking, NeuN primary antibody was added (mouse anti-NeuN monoclonal antibody, 1:1000, Millipore Sigma, MAB377), and incubated for 40 min at 4°C with gentle rocking. The samples were then washed by adding 500 μL of immunostaining buffer and centrifuging at 500 ×g for 5 min at 4°C. Supernatant was discarded and the pellet resuspended in immunostaining buffer containing goat-anti-mouse antibody (Alexa Fluor 647, 1:500) and a nuclear stain (Hoechst 34580; 2,5 μg/ml). Aliquots of unstained, only secondary antibody-treated, and single-stained (Hoechst, NeuN) nuclei were saved for use as controls. The number and integrity of the nuclei were evaluated microscopically after each critical step and before FANS. The typical yield for ∼100 mg of cerebral cortex tissue was between 1−3 × 10^6^ nuclei.

FANS was used to collect two single nuclear suspensions per sample (NeuN^+^ and all nuclei). Sorting was performed using a BD FACSAria II or a Sony SH800. The sheath fluid consisted of PBS with a sheath pressure of 20 psi. Sorting was performed using a 100-μm nozzle tip or microfluidic sorting chip (100-μm). For the excitation of forward scatter (FSC) and side scatter (SSC), a 488-nm laser was employed. Hoechst 34580 and Alexa Fluor 647 were excited using 405-nm and 640-nm lasers, respectively. FANS gating was performed in the following order: FSC height vs. SSC height; SSC area vs. Hoechst fluorescence (bandpass filter 450/50); and Alexa Fluor 647 (bandpass filter 665/30) vs. Hoechst fluorescence. The FSC versus SSC gates were set with permissive limits to discard the smallest and largest particles. Hoechst fluorescence was used to distinguish single nuclei from doublets, clumps, and damaged nuclei. Alexa Fluor 647 was used to distinguish neuronal (NeuN^+^) from non-neuronal nuclei. Controls including unstained, only secondary antibody-treated, and only single primary antibody-treated cell suspensions were included to adjust gates thresholds and minimize false positives from nonspecific staining or autofluorescence. Two populations, all nuclei (Hoechst^+^) and neuronal nuclei (Hoechst^+^/NeuN^+^), were collected. The sorted nuclear suspensions were collected in 1.5-mL Eppendorf tubes containing 100–200 μL of collection buffer consisting of 0.1 M PBS, pH 7.4, and 0.1 U/μL RNase inhibitor. After collection, BSA was added to each tube for a final concentration of 1%. To prevent nuclei from adhering to the tube walls, the collection tubes were precoated with BSA. Precoating was performed by filling the tubes with 10% BSA in PBS for 5 min, followed by rinsing with PBS and drying overnight at 4°C.

### snRNA-seq of postmortem human brain nuclei

We used either a modified Drop-seq method^65^ or the standard 10x Genomics Chromium Single Cell 3’ v2 or v3 assays to profile the transcriptomes of nuclei from postmortem human brain tissue. For Drop-seq, the input single nuclei were diluted to a concentration of 200 nuclei/µl. To encapsulate individual nuclei and barcoded beads (Chemgenes, cat # Makosko-2011-10), we employed a microfluidic system (FlowJEM) and adjusted the flow parameters to generate ∼100 µl (∼0.5 nl) droplets (nuclei loading concentration: 200 nuclei/µl; bead concentration: 165 beads/µl; flow rate: 3 ml/h). With these parameters, both the cell occupancy and the expected doublet rates were ∼5%. These rates were confirmed by observing the beads and Hoechst^+^ nuclei within the droplets by fluorescent microscopy. Standard methods proved challenging for digesting nuclear membranes from human brain nuclei, resulting in low transcript detection. To overcome this, we tested various lysis methods (sarkosyl, SDS, and triton) at different concentrations, with or without heat. Lysis buffers containing 1% sarkosyl yielded optimal results without disrupting droplet generation. Furthermore, brief heating of the droplet-encapsulated nuclei (5 min at 72°C) improved lysis efficiency. Reverse transcription and PCR amplification followed previously described protocols^65^. PCR reactions, each containing 4,000 beads (i.e., 200 nuclei), were individually run and subsequently pooled (typically 5−15 PCR tubes, i.e., 1,000−3000 nuclei) for library preparation and sequencing.

For 10x Genomics, the input single nuclei were centrifuged at 400 ×g for 5 min at 4°C to achieve a concentration of ∼350 nuclei per μL. Nuclear concentrations were determined using a hemocytometer. On average, ∼12,500 nuclei were loaded to capture around 5,000 nuclei per sample (with an expected capture efficiency of ∼40%). cDNA amplification and library construction followed the manufacturer’s instructions.

The paired-end libraries generated by Drop-seq or 10x Genomics were sequenced on either Illumina NextSeq 500 or Novaseq 6000 platforms. A total of 243 samples (184 Drop-seq and 59 10x Genomics) were sequenced in 37 sequencing batches. For each sequencing batch, the concentration of each sample was normalized to the total number of nuclei to ensure similar numbers of reads per nucleus. Nuclei were sequenced to a depth of ∼75,000 reads per nucleus.

### Preprocessing, quality control, and integration of snRNA-seq data

The paired-end raw sequence reads were preprocessed using the Kallisto bustools package (kbpython:0.26.0)^66^. An alignment index was constructed based on the human reference pre-mRNA (GRCh38, Ensembl 105). Following the Lamanno workflow, we generated separate count matrices for spliced and unspliced transcripts. These matrices were then merged to obtain the total nucleus count matrix. The quantification of total transcriptome abundance was performed for each of the three matrices. Downstream analysis, including quality control, integration, cell type annotations, and differential gene expression, was performed using the unspliced transcript counts.

Empty droplets were removed by comparison with ambient RNA levels using the DropletUtils package^67^. The identification of empty droplets was performed by analyzing the knee and inflection points on the cumulative transcript counts plots for each sample individually. Nuclei with an FDR <0.05 were removed, resulting in a total of 665,407 nuclei. Further filtering was applied to exclude nuclei with fewer than 200 genes, leaving 549,074 nuclei.

To identify potential doublets, we used the DoubletFinder package version 4.2^68^. Among the 10x Genomics samples, an average doublet rate of 2.85% and 1.74% was detected in v2 and v3 samples, respectively, while the Dropseq samples had a doublet rate of 0.003%. The identified doublets were labeled and retained during batch correction and data integration. Following clustering and dataset annotation, the majority of labeled doublets clustered together. These clusters, containing doublets, were excluded from further downstream analysis.

To analyze the raw count data, we used Scanpy in the python package version 3.9.1^69^. First, we used a series of preprocessing steps for normalization and scaling. Highly variable genes were identified using default parameters and a dispersion threshold of 0.5. Principal Component Analysis (PCA) was applied to reduce dimensionality, generating 50 principal components. Subsequently, a neighborhood graph was constructed using default parameters with 15 neighbors, and Leiden graph-based clustering was performed using correlation distance metrics. To address batch effects and integrate data from different brain regions and disease stages, we evaluated various methods for batch correction and integration, including SCT, Harmony, BBKNN, and Scvi-tools. We selected Harmony (v1.2.2769)^70^ as the integration method based on silhouette score values of 0.8 or higher, as well as visual inspection of UMAP plots representing experimental assay, sequencing batch, donor, brain region, disease stage, sex, and UMI abundance. The selected integration variables were the experimental assay (Dropseq, 10x Genomics v2, and 10x Genomics v3) along with brain region (BA9, BA7, and BA17). After integration, the neighborhood graph and Leiden graph-based clustering were generated again. Marker genes for each cluster were determined using the Wilcoxon rank sum test with a significance threshold of adjusted p-value (padj) < 0.05.

The integrated dataset contained 549,074 nuclei. To further optimize the lower gene cutoff, thresholds from 200 to 500 genes were tested, and 300 genes were chosen as the final cutoff. All clusters comprised nuclei from every donor, and no clusters exclusively contained nuclei with low UMI counts. Three small clusters containing doublets, totaling 17,442 nuclei, were excluded. Mitochondrial gene content was measured and annotated, but only outlier nuclei with higher than 5% mitochondrial genes (1,778 nuclei) were discarded, resulting in a total of 424,528 high-quality nuclei (362,224 neuronal and 62,304 non-neuronal) for downstream analysis.

### Major cell type, neuronal subtype, and glial cell state annotations

The major neuronal and non-neuronal populations were identified based on the expression of known marker genes: *SLC17A7* (excitatory neuron), *GAD1* (inhibitory neuron), *FGFR3*, *AQP4*, and GFAP (astrocyte), *CSF1R*, *CX3CR1*, and *CD163* (microglia), *PLP1* and *MOG* (oligodendrocyte), *PDGFRA* and *CSPG4* (OPC), *CLDN5* and *FLT1* (endothelial), *NOTCH3* (pericyte), and *CYP1B1* and *COL15A1* (VLMC).

These major cell type clusters were subsetted and reclustered within the integrated PC space to identify neuronal subtypes and glial states. Clustering reliability was determined based on silhouette score values of 0.8 or higher and WCSS (Within Cluster Sum of Squares). The first 30 PCs and a resolution of 1.0 were employed for both the excitatory subset (282,930 nuclei) and the inhibitory subset (79,294 nuclei). Marker genes for each cluster were ranked using the Wilcoxon rank sum test with the following criteria: minimum expression fraction (either in the tested cluster or in all other nuclei combined) of 0.2, log-fold change > 0.5, padj < 0.05. After merging two excitatory clusters that lacked marker genes to reliably distinguish between them and discarding two small inhibitory clusters (207 nuclei) with mixed markers, a total of 18 excitatory (Ex) and 19 inhibitory (In) clusters were obtained. We visualized the UMAP with a minimum distance of 0.6 and a spread of 1.4.

To identify the top marker genes for each cluster, the following criteria were applied: expression fraction within the cluster (pts) > 0.2; expression fraction within all other nuclei (pts_rest) < 0.1; ratio pts/pts_rest > 3; log-fold change > 1.5; and padj < 0.05. For Ex1, Ex2, and Ex5, the pts_rest was set at < 0.2 and the ratio pts/pts_rest was set at > 2. The clusters were named based on canonical markers for major subclasses (i.e., *CUX2*, *RORB*, *THEMIS*, and *FEZF2* for excitatory; and *LHX6* and *ADARB2* for inhibitory), followed by 1−3 top marker genes. Additionally, we compiled gene sets consisting of 7-10 genes for each neuronal subtype, selected from the top marker genes. These marker genes and gene sets precisely labeled each of our annotated neuronal subtypes in our dataset and a reference dataset^23^, and thus are useful to identify neuronal subtypes computationally and by histology.

To compare our neuronal clusters and their marker genes with a reference dataset, we utilized a publicly available dataset containing over one million neuronal nuclei from the DLPFC of donors with dementia and healthy controls^23^. To determine the degree of similarity between the annotations in the two datasets, we subset both count matrices keeping only highly variable genes (3,000 genes), identified the top 10 markers for each cluster, and calculated the cosine similarity distance between the mean expression values of genes for each cluster. A lower distance in the similarity matrix indicates a higher level of agreement.

The non-neuronal subset clusters included: astrocytes (14,691), microglia (5,071), oligodendrocytes (36,589), OPCs (5,770), and vascular cells (183). These populations were reclustered using the first 10 PCs, with a resolution of 0.3 for astrocytes, 0.2 for microglia and oligodendrocytes, and 0.1 for other types. The top marker genes for each cluster were identified using the same method as for the neuronal clusters, using the following thresholds: pts > 0.2; pts_rest < 0.1; ratio of pts/pts_rest > 2.5; log-fold change > 1.0; and padj < 0.05. We annotated 4 astrocyte, 4 microglia, and 2 oligodendrocyte cell states. Other non-neuronal types were not subclassified further due to their relatively low nucleus numbers.

### Cell identity prediction using scANVI

We employed the scANVI^71^ machine learning method to predict the identity of unannotated neuronal nuclei with relatively low UMI counts from our dataset and to predict the identity of neuronal populations from public datasets using the annotations from our dataset. For model training, we utilized our excitatory (18 neuronal clusters) and inhibitory (19 neuronal clusters) datasets as training set. We selected 2,000 highly variable genes and the top 200 marker genes for each cluster (Wilcoxon rank sum test, log-fold change > 0.8, padj < 0.05), along with our cell-type-specific gene sets. The model underwent training for a maximum 200 epochs with 3 layers and 50 latent spaces. To address batch effects during training, we introduced a combined batch effect key considering both the profiling assay (DropSeq, 10x v2, 10x v3) and brain region (BA9, BA7, and BA17). We monitored model convergence and loss for each epoch using an elbow plot to determine the optimal number of epochs for effective convergence. To prevent overfitting, we employed a single-layer perceptron with the ‘linear_classifier’ parameter set to ‘True’, promoting model simplicity and reducing bias towards the training data. Additionally, we applied the var activation (torch.nn.functional.softplus) function to ensure stable optimization. To represent rare cell types adequately, we set the ‘n_samples_per_label’ to 1000. Probabilities of cell cluster assignments from the latent space were computed using the ’soft’ function, providing a confidence measure for each prediction. Model accuracy was assessed by comparing true labels with predictions using sensitivity and specificity measures. Additionally, we conducted differential testing using the Wilcoxon rank sum test, employing predicted annotations to validate the reliability of the neuronal subtypes inferred from our predictions. We assigned identity to query data using a probability threshold of 0.99 to minimize false positives. We predicted excitatory cell identity labels for five public reference datasets (SEA-AD DLPFC and MTG; Mathys *et al*., 2023; Green *et al*., 2024; Jorstad *et al*. 2023)^3, 5, 8, 23^ to demonstrate that the pretrained model can be applied across datasets.

### Neuronal subtype proportion quantification

To quantify the relative proportions of each neuronal subtype within each disease group (low, intermediate, high pathology) and each region (BA9, BA7, BA17), we calculated the relative abundance of each subtype per donor in relation to all neurons and either excitatory or inhibitory neurons. To test for statistically significant differences in cell composition among disease groups, we conducted three different analyses: beta regression (modeled for data measured as proportions), Kruskal-Wallis H test (a rank-based nonparametric test for non-normally distributed data), and a linear mixed model (for modeling variances and covariances). Each of the three analyses was conducted both with and without the inclusion of nuclei with relatively low UMI counts (200-300 genes per nucleus), as annotated by scANVI predictions.

For beta regression, we utilized the R package betareg version 3.1-4 with the formula Relative.Abundance ∼ Condition and the bias-corrected maximum likelihood estimator (R > betareg(Relative.Abundance ∼ Condition, data=data, type=“BC”)). We excluded data from two donors for BA9 and one donor for BA17 from this analysis due to their low cell counts (<500 nuclei). To account for multiple hypothesis testing, we applied Holm’s method to adjust the p-values obtained from beta regression using the ‘p.adjust’ function of the R Stats package, considering the number of tested neuronal subtypes as the ’length’ variable.

We conducted linear mixed modeling using both absolute cell counts and cell proportions. The fixed covariate was profiling assay (Dropseq, 10x v2, and 10x v3). Donor was considered a random effect. Our analysis focused on evaluating the differences in each neuronal subtype across the disease group (used as a predictor), with the low pathology group serving as the reference category. Subsequently, we computed the coefficients, t-values, p-values, and corresponding Z-scores for each neuronal subtype, and applied the Benjamin-Hochberg correction to account for false positives in the p-values.

Differential population analysis was performed using scCODA^34^, which applies a Bayesian model to assess significant changes in cell populations. We generated raw count data (donor * Author_Annotation) for each brain region (BA9 and BA17), with donor as the covariate. We then calculated cell type proportions, fitted the Bayesian model to the data, and evaluated the effects across disease progression groups.

### Spatial transcriptomics

We used the 10x Genomics Visium platform (Spatial 3’ v1 chemistry) to spatially map 37 neuronal subtypes (18 excitatory and 19 inhibitory), astrocytes, oligodendrocytes, OPCs, and microglia in fresh-frozen tissue sections from the neocortex of AD and healthy control donors. A total of 16 tissue sections were studied, including controls (BA9, BA7, BA17, motor cortex, entorhinal cortex, and hippocampus from a control donor, and two additional sections of BA9 and BA7 from another donor) as well as AD (BA9 and BA7 from 3 donors with high AD pathology and BA9 from 2 donors with intermediate pathology). The sections were cut at a thickness of 12 µm on a cryostat and mounted on fiduciary frames of four 10x slides. H&E staining was performed, and the slides were temporarily coverslipped with mounting medium (85% glycerol containing 0.2 U/μL RNAse inhibitor) and digitally scanned at 200x magnification using a Zeiss Axio Imager M2 microscope equipped with a color digital camera (Axiocam) and MBF Stereo Investigator with a 2D slide scanning extension module. Permeabilization enzyme treatment was applied to the tissue for 15 minutes at 37°C, as determined by the Tissue Optimization protocol provided by 10x Visium. Reverse transcription, second strand synthesis, and cDNA amplification were carried out according to the manufacturer’s recommendations. We utilized the targeted Human Neuroscience gene expression panel, which consists of 1,186 genes, and supplemented it with a custom panel comprising 197 cell type-specific marker genes. The marker genes were selected from our snRNA-seq dataset based on their specificity to label our annotated neuronal clusters and their expression levels. The custom hybridization capture panel oligos were obtained from IDT (IDT NGS Discovery Pools). Library sequencing was performed on the Illumina Novaseq 6000 platform at a depth of 15,000 reads per spot, resulting in a sequencing saturation of approximately 96%. This 1,383 gene panel provides a cost-efficient tool for mapping neuronal vulnerability in the human AD brain while allowing to sequence at a 90% lower cost compared to whole transcriptome sequencing. To evaluate the quality of our spatial data, we used the 10x Space Ranger pipeline, which maps the transcriptomic data on the high-resolution microscopic images. On average, we detected 1,336 out of the total 1,383 targeted genes. The median number of targeted genes and UMI counts detected per spot were 221 and 392, respectively.

### Integration of snRNA-seq and spatial transcriptomics

We used Stereoscope^72^ to integrate the spatial transcriptomics (ST) data generated with 10x Genomics Visium and the snRNA-seq data. First, we subsetted the snRNA-seq data based on the genes present in the Visium spatial transcriptomics dataset (1,383 genes). We trained variation auto encoder model using the snRNA-seq data to construct a single-cell reference latent variable for inferring cell type-specific gene expression patterns. For model training, we used the following parameters: layer = ‘unspliced’; labels_key = ‘Author_Annotation’; max epochs = 200. We checked the model convergence by elbow plot. Then, we trained the spatial model using the Visium data and the pre-trained snRNA-seq data for a maximum of 2000 epochs. This allowed us to identify cell types using negative binomial latent variables. We performed different iterations to visualize cell populations in the Visium space considering different combinations of cell types (i.e., all major cell types; each of the excitatory neurons, inhibitory neurons, and glial populations; and each excitatory neuronal subtype). To visualize each of the cell subtypes in each ST tissue section slide, we utilized the matplotlib and seaborn plotting Python packages.

### Co-expression network analysis

We performed weighted gene co-expression network analysis (WGCNA) on our high-dimensional snRNA-seq data using the hdWGCNA R package^73^ to compute co-expression gene modules of interconnected genes within each neuronal cell subtype and brain region. We constructed meta-cells using the following parameters: group.by = “Author_Annotation”, k = 25, and minimum cell threshold of 50. After constructing the meta-cells, we normalized the object using default parameters, including the use of Harmony for dimensional reduction and batch correction. We then constructed co-expression networks. Gene correlations were transformed into a similarity matrix using the power function, which preserves strong correlations. Modules were identified through hierarchical clustering with similarity distance measures. Module reliability and robustness were estimated using bootstrap resampling with 5,000 iterations. We ranked the highly correlated genes, defined as the kME (module eigengene), by their kME values for each neuronal cell subtype within the modules and retained the top 50 intra-module co-expressed genes.

### Differential gene expression analysis

We performed differential gene expression (DGE) analysis to compare the low vs intermediate pathology groups (designated as ‘early’ changes) and the intermediate vs high pathology groups (designated as ‘late’ changes), within each neuronal subtype and brain region. To ensure the reliability of our analysis, we employed various methods, including a zero-inflated regression mixed model implemented in MAST^74^ and lme4^75^, bootstrap resampling with 100 iterations, and pyDESeq2^76^ on pseudobulk aggregated counts.

For the zero-inflated regression mixed model (MAST and lme4), we used the following model formula:

Zlm(∼ condition + (1 | donor) + cngeneson + Assay + Age + Sex + RIN + total_counts, sca, method = ’glmer’)

In this model, donor is considered a random effect. The fixed covariates include “cngeneson“ (i.e., cellular gene detection rate), age, sex, RIN, and total raw sequencing counts. The DE genes were filtered using the following thresholds: percentage of expression > 20% for at least one condition, ∣logFC∣>0.1, and false discovery rate < 0.05.

To ensure the robust and reproducible identification of significant DE genes, we employed bootstrapping followed by DGE (MAST/lme4, as detailed above). This resampling technique mitigates potential effects from outliers and the varying number of nuclei per cluster. We conducted a series of 100 iterations. In each iteration, we randomly selected 50% of nuclei from each neuronal cell subtype from each comparison. Subsequently, we computed the DE genes for each iteration and assigned confidence scores based on frequency analysis across all iterations. We filtered the DE genes based on their consistent identification as differentially expressed, retaining those genes that exhibited the same significant up or downregulation in at least 20 out of the 100 iterations.

Additionally, we used a pseudobulk aggregation method with raw gene abundance counts to construct representative expression profiles for each neuronal subtype within each donor, disease group, and brain region. The data were organized with donors as rows and genes as columns. Next, we aggregated the data from individual donors into a single pseudobulk count dataset. We then performed log normalization on the raw count data and applied a gene filter, retaining only genes expressed in at least 20 nuclei. Subsequently, we created a DESeq2 object using pyDESeq2. To evaluate the data’s inherent variability, we conducted principal component analysis (PCA), a high-dimensional reduction technique. For DGE analysis, we established a design matrix to compare disease groups. To ensure statistical significance, we applied the Benjamin-Hochberg correction method with a threshold of padj < 0.05.

We defined ‘high-confidence’ DE genes as those identified by at least two different methods: the mixed model and either one of the other DGE methods (bootstrap or pseudobulk) or network co-expression analysis (top 50 genes by kME values from the hdWGCNA). This comprehensive approach aimed to enhance the reliability of our DGE predictions by ensuring consistency across methods and experimental conditions.

### Functional enrichment analysis

We used multiple methods for functional enrichment analysis, including Enrichr (Python), Metascape^77^, and g:profiler^78^. The input data consisted of high-confidence DE genes obtained from comparing low vs. intermediate and intermediate vs. high pathology groups (’early’ and ’late’ changes) within each neuronal subtype and brain region. For Enrichr^79^, we used brain-specific gene sets from the Genotype-Tissue Expression(GTEx)^80^ and Synaptic Gene Ontology (SynGO)^81^ databases to establish background gene expression. Statistical significance thresholds were determined using an adjusted p-value < 0.05 and at least a minimum of three genes per group. We utilized Metascape with the following custom parameters: a minimum overlap of 5, a p-value threshold of 0.01, and a minimum enrichment score of 2.5. The top 50 enriched functional modules were visualized in heatmaps using the Matplotlib and Seaborn Python packages.

Additionally, we used g:profiler to perform functional enrichment analysis for Ex2 L2/3 IT and Ex5 L4 IT, using as input the top 50 co-expressed network genes from each module from our hdWGCNA analysis. We selected key driver GO terms within the Molecular Function and Biological Process categories, with GO terms having a size between 10 and 1000 and an adjusted p-value threshold of 0.05. The top enrichment terms from each module were visualized in a dot plot using Matplotlib and Seaborn, and the enrichment networks were visualized using Cytoscape^82^.

### Immunohistochemistry in human brain tissue

Immunofluorescence to quantify KCNIP4 was performed on 20 µm-thick cryosections of fresh-frozen brain tissue. Sections were fixed with 2% PFA for 20 minutes, blocked with 10% normal goat serum (NGS) and 2% BSA in PBS with 0.25% Triton X-100 (PBT) for 1 hour at RT, and then incubated overnight at 4°C with primary antibodies in PBT containing 3% NGS and 0.5% BSA. The primary antibodies used were guinea pig anti-NeuN (1:50, Sigma ABN90), rabbit anti-EYA4 (1:50, Thermo Fisher Scientific PA552113), and rabbit anti-KCNIP4 (1:50, Proteintech 60133-1-Ig). After three washes with PBT, sections were incubated with secondary antibodies for 2 hours at RT: Alexa Fluor 647 anti-guinea pig (Invitrogen, A21450; 1:200), Alexa Fluor 488 anti-rabbit (1:200, Invitrogen A11070;), and Alexa Fluor 546 anti-mouse (Invitrogen, A11018; 1:200). Sections were counterstained with DAPI (1:2000, Invitrogen) for 20 minutes, rinsed in PBS, mounted with aqueous mounting medium (Invitrogen P36930), and sealed. Images were acquired using a Zeiss LSM980 laser scanning confocal microscope with consistent parameters and processed with CellProfiler using custom pipelines for automatic cell segmentation based on NeuN and analysis of EYA4-positive cells and KCNIP4 intensity.

Immunohistochemistry for VGLUT2 and NeuN was performed on 50 µm-thick free-floating fixed sections, obtained from tissue blocks fixed with 4% PFA for 3 days, cryoprotected in 30% sucrose, and sectioned on a sliding microtome. The free-floating sections were rinsed in PBT and incubated in PBT containing 1% hydrogen peroxide for 30 minutes to block endogenous peroxidase activity. After washing with PBT, sections were incubated in a blocking solution containing 10% NGS and 2% BSA for 1 hour at RT. Sections were then incubated with mouse anti-NeuN (1:1000, Millipore Sigma MAB377) or mouse anti-VGLUT2 (1:1000, Millipore Sigma MAB5504) diluted in PBT containing 3% NGS and 0.5% BSA overnight at 4°C. After washing, sections were incubated with a biotinylated goat anti-mouse antibody (VectorLabs BA-9200) diluted in PBT containing 3% NGS for 2 hours at RT, and then washed again. This was followed by incubation with ABC solution (Vectastain Elite ABC-HRP kit, VectorLabs PK-6100) for 1 hour at RT. For the chromogenic reaction, 3,3’-Diaminobenzidine (DAB) substrate solution (Sigma D5905) was used. Sections were air-dried, dehydrated with ethanol followed by xylene, and coverslipped with Permount mounting medium.

### Experimental animals

*App* knock-in mice (B6.Cg-*App^tm^*^1^*^.1Dnli^*/J, strain #034711; also known as *App*^SAA^) and WT controls (C57BL/6J, strain #000664) were obtained from The Jackson Laboratory (Bar Harbor, ME, USA). The mice were bred on a pure BL6/J background, and genotypes were confirmed by real time PCR (Transnetyx, Cordova, TN). All mice were housed in a barrier facility and maintained according to the guidelines for animal testing and research under a protocol approved by the Stanford’s Administrative Panel on Laboratory Animal Care (APLAC). Previous studies have reported sex differences in some behavioral assays, but not in pathology^24^. Due to the small sample size, we used only male mice in this study.

### AAV-driven KCNIP4 expression in excitatory neurons in adult mice

Twelve-month-old *App^SAA^* and control male mice were injected retroorbitally with 1 × 10¹¹ vg of AAV-PHP.eB-CaMKIIa-*Kcnip4*-P2A-EGFP in 100 µL of PBS or with AAV-PHP.eB-CaMKIIa-EGFP as a control. Cloning and AAV production were performed by Vector Biolabs (mouse *Kcnip4* isoform 1; NCBI Reference Sequence: NP_001186171.1). Thirty days post-injection, mice were perfused with 0.9% saline for 3 minutes, and the brains were extracted. The right hemisphere was frozen in isopentane at -50°C, while the left hemisphere was fixed in 4% PFA for 24 hours, followed by cryoprotection in 30% sucrose for 24 hours. The fixed tissue was cut into 50 µm-thick free-floating sections and stored at - 20°C in a cryoprotectant solution (30% glycerol, 30% ethylene glycol in PBS) until further processing.

### Immunoblotting of mouse cortex tissue

Frozen mouse brain cortex was lysed in tris/SDS/glycerol buffer, and protein concentration was determined using a BCA assay (Thermo Fisher Scientific). Twenty μg of protein were separated on 10% Mini-PROTEAN® TGX™ Precast Protein Gels (Bio-Rad) and transferred to PVDF membranes using the Trans-Blot Turbo (Bio-Rad) semi-dry transfer system. The membranes were blocked with 5% milk in tris-buffered saline with 0.05% Tween 20 (TBS-T) for 60 minutes at RT and then incubated overnight at 4°C with mouse anti-Kcnip4 antibody (1:1,000, Proteintech 60133-1-Ig) and anti-GAPDH antibody (1:10,000, Thermo Fisher Scientific MA5-15738). After three 10-minute washes with TBS-T, the membranes were incubated with a goat anti-mouse secondary antibody (1:1,000, Invitrogen G-21040) for 60 minutes at RT. Following washing with TBS-T, membranes were developed using ECL (Thermo Fisher Scientific, 32106) and imaged on X-ray film (Thermo Fisher Scientific, 34091). Images were processed and quantified using ImageJ.

### Immunohistochemistry in mouse brain tissue

Immunohistochemistry was performed on 50-µm-thick free-floating slices of mouse brain tissue. For immunofluorescence, slices stored in cryoprotectant solution were washed three times for 10 minutes each with PBS, then photobleached under full spectrum LED light for 48 hours in a cold chamber^4^. The sections were blocked with 10% NGS and 2% BSA in PBT for 1 hour at RT and incubated overnight at 4°C with primary antibodies in PBT containing 3% NGS and 0.5% BSA. For c-Fos and Arc quantification, we used: guinea pig anti-c-Fos (1:200, Synaptic Systems 226 308), mouse anti-Arc (1:200, Synaptic Systems 156 111), rabbit anti-GFP (1:1000, Invitrogen A11122), and anti-NeuN (1:200, Millipore Sigma ABN90P). After primary antibody incubation, sections were washed three times for 15 minutes each with PBT and then incubated with secondary antibodies (1:200, Alexa Fluor 647 anti-guinea pig, Invitrogen A21450; Alexa Fluor 488 anti-rabbit, Invitrogen A11070) for 2 hours at RT. Following three additional 15-minute washes, tissues were counterstained with DAPI (1:2000, Invitrogen) for 30 minutes at RT. The slices were rinsed in 0.05M TBS, mounted with aqueous mounting medium (P36930, Invitrogen), and sealed. Images were acquired using a Zeiss LSM980 laser scanning confocal microscope with consistent parameters across all samples. Image processing was carried out using CellProfiler with custom pipelines for automatic segmentation of GFP**^+^** and GFP**^−^** neurons based on NeuN and GFP markers, and quantification of c-Fos and Arc.

For chromogenic immunohistochemistry, free-floating sections were incubated in 0.6% hydrogen peroxide in PBT for 20 minutes to block endogenous peroxidase activity. The primary antibodies used included rabbit anti-human amyloid beta (1:500, IBL 18584), rabbit anti-GFAP (1:2000, Dako Z0334), and rabbit anti-Iba1 (1:500, FujiFilm 019-19741). Sections were then incubated with a secondary antibody, followed by an avidin/biotin-based peroxidase system and chromogenic detection using DAB, as previously described for human tissue. Brightfield images were captured with a Zeiss Axio Imager 2 and a Hamamatsu digital camera (C11440), and the stained cortical area was quantified using ImageJ.

## Data availability

The raw snRNA-seq data, associated metadata, and processed digital expression matrices have been deposited at the NCBI’s Gene Expression Omnibus with accession number GSE263468. Eight of 243 samples were included in previous studies (GSE129308 and GSE181715). The datasets are publicly available for interactive viewing and exploration on the cellxgene platform at https://cellxgene.cziscience.com/collections/0d35c0fd-ef0b-4b70-bce6-645a4660e5fa

## Acknowledgements

Human tissue was obtained from Stanford’s Department of Pathology and Alzheimer’s Disease Research Center (NIH/NIA P30AG066515), UCLA Department of Pathology and Easton Center, and the NIH Neurobiobank (Sepulveda repository in Los Angeles, CA, and Mt. Sinai Brain Bank in New York City, NY). This work was supported by grants to I.C. from NIH/NIA (R01AG059848, R01AG08214702), BrightFocus (A20173465), the Alzheimer’s Association (AARG-17-528298), and the Chan Zuckerberg Initiative (Ben Barres Early Career Acceleration Award, grant ID 199150).

## Author contributions

Conceptualization: S.A.P.D., I.C.; Human tissue procurement and Neuropathology: K.V., I.C.; Single-nuclear and spatial transcriptomics data generation: M.O.G., J.P.; Data analysis: S.A.P.D., W.T.; Histology: J.S.R., J.P., K.V.; Functional assays in mice: J.S.R.; Funding acquisition: I.C.; Supervision: I.C.; Writing: S.A.P.D., J.S.R., I.C.; All authors read and approved the final manuscript.

## Ethics declarations

### Competing interests

The authors declare no competing interests.

## Supplementary material

**Supplementary Table 1**. Brain tissue samples and quality control (QC).

**Supplementary Table 2**. Gene markers for excitatory neurons (Ex 1-18), interneurons (In 1-19), and glial cell states, and gene sets (7-10 cluster-defining genes) for each cell subtype. Score = Wilcoxon rank-sum test variance score (Z score); pts = fraction of nuclei in the cluster expressing the gene; pts_rest = fraction of nuclei in all other clusters expressing the gene; Cutoff applied: pts > 0.2; pts_rest < 0.1 (< 0.2 for Ex1, Ex2, Ex5); padj < 0.05.

**Supplementary Table 3**. Estimated numbers of nuclei within each neuronal cluster (absolute numbers and percentage of nuclei relative to total neurons or to either excitatory or inhibitory types) for each donor across regions and disease stages.

**Supplementary Table 4**. ’High-confidence’ DE genes in excitatory neuron clusters across brain regions and disease stages. ’High-confidence’ DE genes were defined as those identified by at least two methods: a mixed model (implemented in MAST and lme4) and either another DGE method (bootstrap or pseudobulk) or network co-expression analysis (top 50 genes by kME values from the hdWGCNA).

**Supplementary Table 5**. Gene ontology and pathway enrichment analysis of ’high-confidence’ DE genes using Metascape with custom parameters: a minimum overlap of 5, a p-value threshold of 0.01, and a minimum enrichment score of 2.5.

**Supplementary Table 6**. ’High-confidence’ DE genes in clusters representing either vulnerable (Ex2; L2/3 IT) or resistant (Ex5; L4 IT) neuronal subtypes in BA9 and BA17 during early and late disease stages.

**Supplementary Table 7**. Co-expression modules identified through hdWGCNA for each excitatory cluster in BA9 and BA17.

---

**Extended Data Fig. 1.**
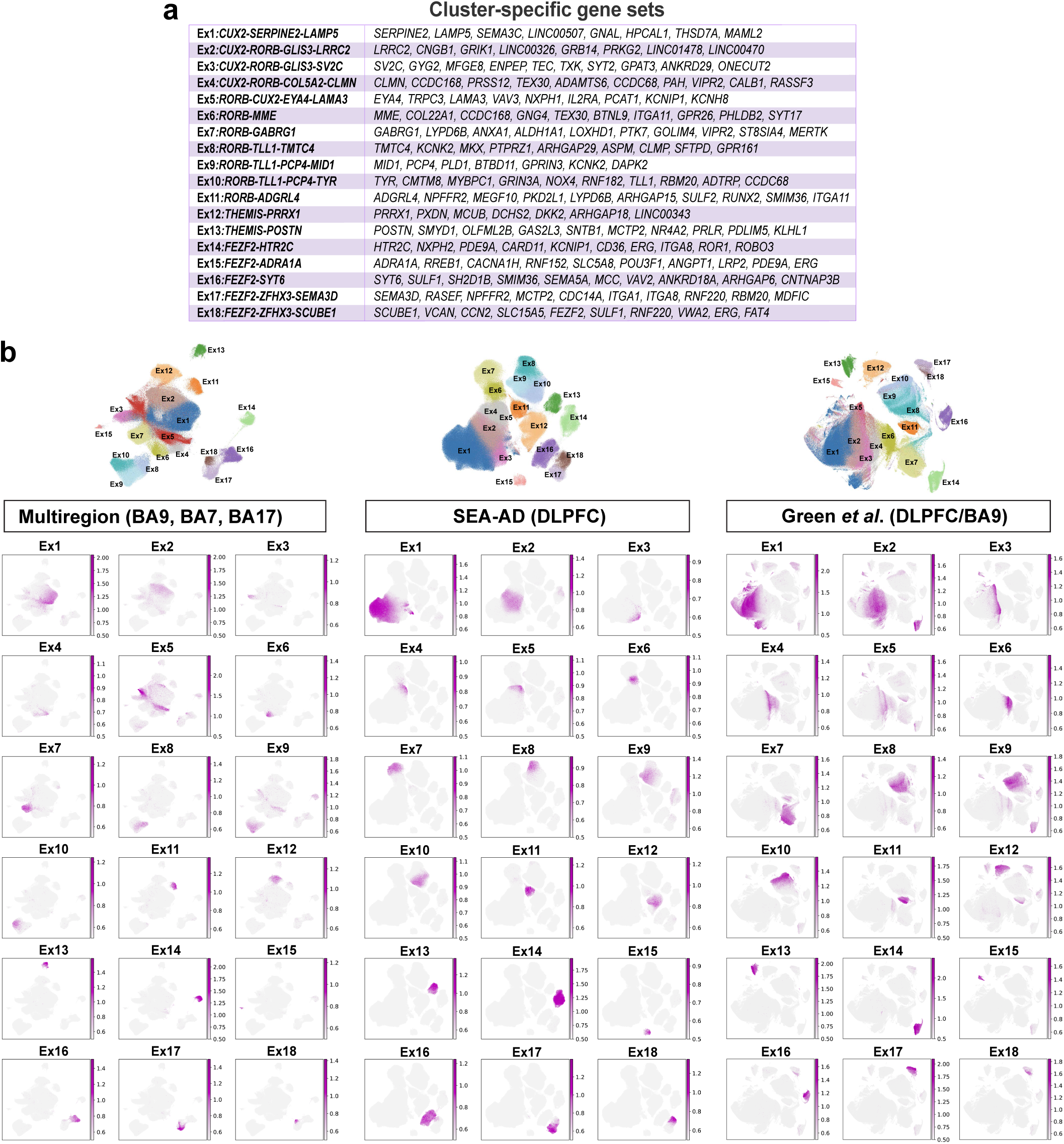
Identification of excitatory neuron clusters across datasets. **a**, Cluster-defining gene sets for each excitatory neuron cluster (Ex1−18). **b**, UMAP visualization of the excitatory neuron clusters (top) and gene expression UMAPs for the genes included in the cluster-specific gene sets (bottom), across three datasets: the present manuscript (multiregion; BA9, BA7, BA17), SEA-AD (DLPFC)^23^, and Green and colleagues (DLPFC/BA9)^5^. Clusters in the public datasets were predicted using scANVI.

**Extended Data Fig. 2.**
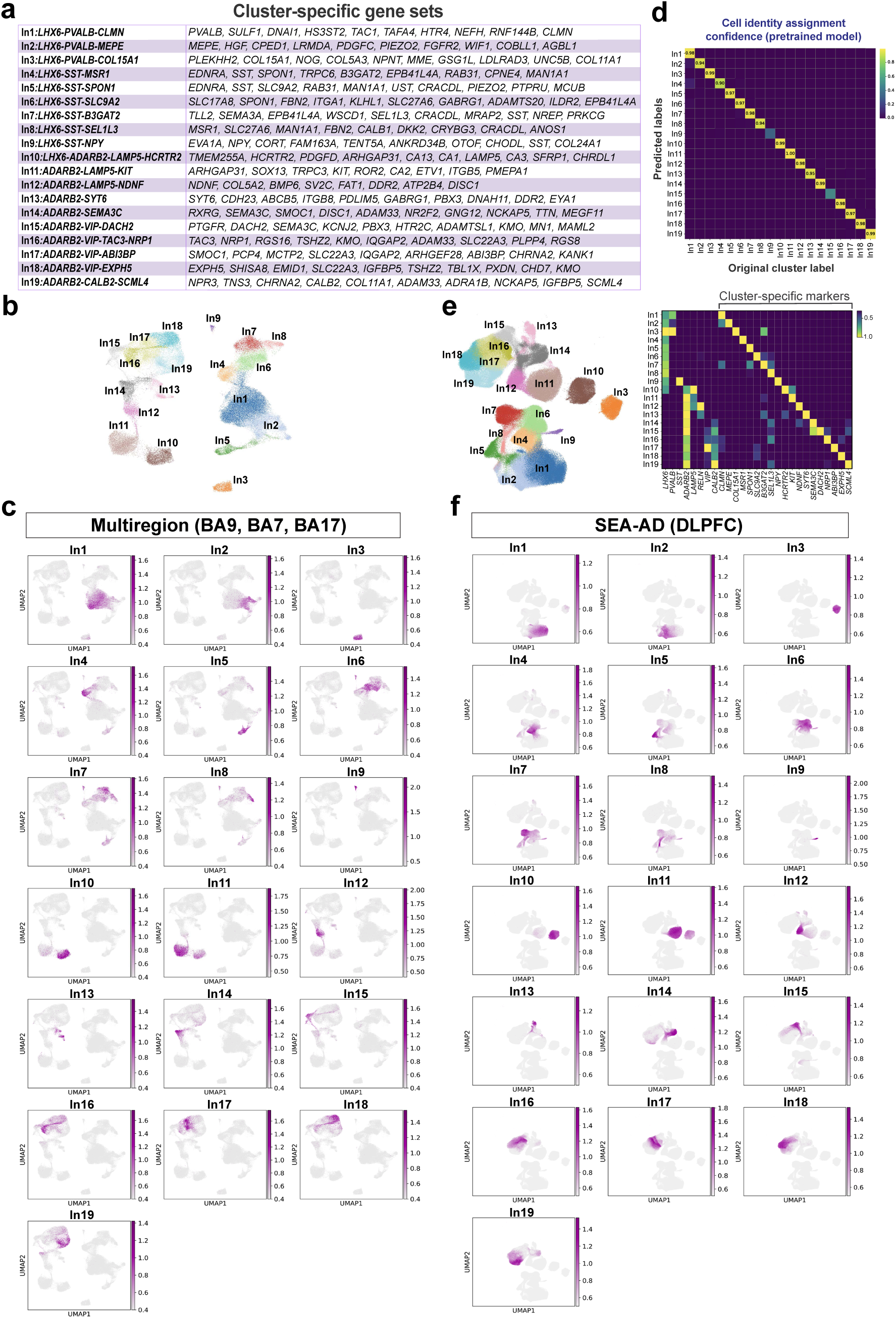
Identification of inhibitory neuron clusters across datasets. **a**, Cluster-defining gene sets for each inhibitory neuron cluster (In1−19). **b**, UMAP visualization of the inhibitory neuron clusters **c**, UMAP visualization of gene expression for the genes included in the cluster-defining gene sets. **d**, Heatmap showing the assignment confidence scores for each inhibitory cluster from the pretrained model using scANVI. **e**, UMAP and heatmap showing the predicted inhibitory clusters in a reference, publicly available dataset from the prefrontal cortex (SEA-AD DLPFC^23^) and their marker genes. **f**, UMAP visualization of gene expression for the cluster-defining gene sets in the SEA-AD (DLPFC)^23^ dataset. Clusters in the public dataset were predicted using scANVI.

**Extended Data Fig. 3.**
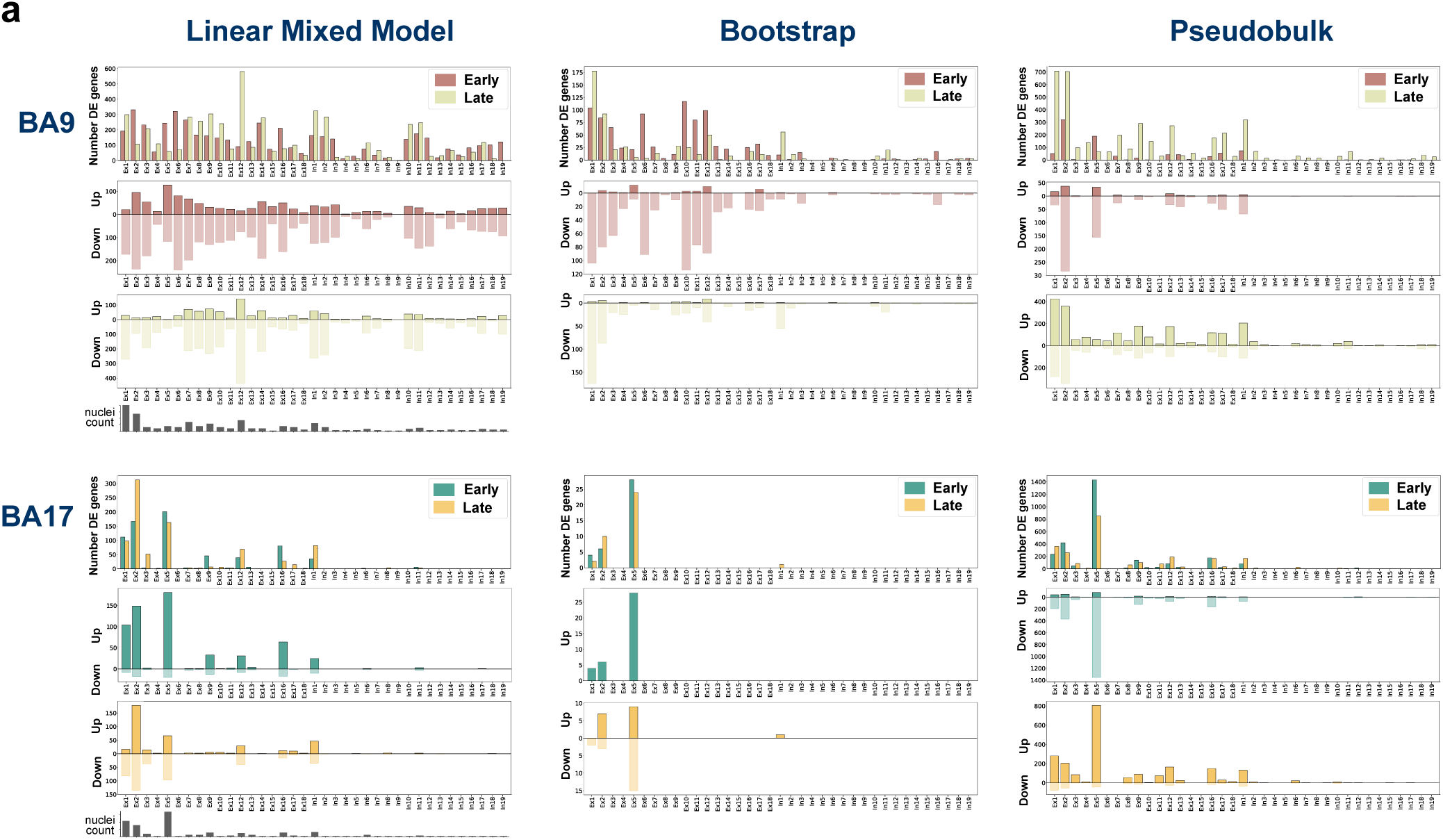
Number of DE genes identified using a linear mixed model, bootstrap, or pseudobulk. **a**, Bar plots illustrate the total numbers of DE genes, upregulated genes, and downregulated genes within each excitatory and inhibitory neuronal cluster at early and late disease stages in the BA9 and BA17. The number of nuclei per cluster is plotted for reference.

## Notes

### Competing Interest Statement

The authors have declared no competing interest.

